# Adaptation of pain-related projection neurons in acute but not chronic pain

**DOI:** 10.1101/2024.05.06.592712

**Authors:** Ben Title, Enrique Velasco, Nurit Engelmayer, Prudhvi Raj Rayi, Roy Yanai, Shmuel Hart, Ben Katz, Shaya Lev, Yosef Yarom, Alexander M Binshtok

## Abstract

Pain hypersensitivity is associated with increased activity of peripheral and central neurons along the pain neuroaxis^1^. On the other hand, in other neuronal systems, increased activity leads to adaptive reduction of neuronal excitability to maintain homeostasis^2–4^. Projection neurons (PNs) of spinal and medullary dorsal horns summate the activity of primary nociceptive and local central interneurons and convey it to higher centers^5^. We show that at the peak of acute inflammatory pain, PNs reduce their intrinsic excitability and, consequently, action potential firing. When pain resolves, the excitability of PNs returns to baseline. Using electrophysiological and computational approaches, we found that an increase in potassium A-current (I_A_) underlies the decrease in the excitability of PNs in acute pain conditions. We hypothesized that an I_A_-induced decrease in PNs firing may restrain the output from the dorsal horn to prevent sensitization and pain chronification. Indeed, no changes of I_A_ in PNs were observed in chronic pain conditions, and PNs exhibit increased intrinsic excitability and firing. Our results reveal an adaptive mechanism in acute pain conditions for regulating the output from the dorsal horn network, which, if interrupted, could trigger pain chronification.

## Introduction

Controlled, coordinated, and time-limited avoidance responses to potentially harmful stimuli are imperative for survival. These avoidance responses are governed by a nociceptive system tuned to detect and respond to injury-inducing stimuli^6–8^. Noxious, injury-inducing stimuli are detected and encoded by primary nociceptive neurons^9–12^ that convey the information into the superficial laminae of the spinal or medullary dorsal horn^7^. There, the information is processed by the intricate network of interneurons^13, 14^ and relayed to higher centers via projection neurons (PNs)^15^, which integrate the inputs from primary afferents and dorsal horn interneurons^5^. In pathological conditions, such as inflammation or nerve injury, primary afferents become hyperexcitable, thus modifying the rate and timing of their firing toward the nociceptive dorsal horn network^9, 16–19^. There, it produces overall increased activity of dorsal horn interneurons^20–22^, resulting in amplified activation of PNs^23–25^, thus enhancing the output of the first CNS nociceptive networks to higher centers and leading to increased nociception and pain^25–28^.

Here, we asked whether and how changes in PNs properties contribute to the enhanced output of the dorsal horn in pathological conditions. Strikingly, we show that in acute pain conditions, PNs undergo an adaptive decrease in intrinsic excitability and action potential firing when the hyperalgesia and pain peak. The excitability of the PNs returns to baseline values when mice recover from hyperalgesia and pain. Further, we demonstrate that the decreased AP firing of PNs in acute pain conditions was mediated by the increase of potassium A-current (I_A_). In contrast, in chronic pain conditions, no modulation of I_A_ occurred, and PNs showed a substantial increase in excitability.

In other neuronal systems, it has been shown that enhanced synaptic input or neuronal activity causes adaptive changes in intrinsic excitable properties, reducing neuronal excitability, hence regulating neuronal homeostasis and tuning down neuronal output^2, 4, 29^. Our results suggest that during tissue injury or inflammation, the output of pain-related PNs is refined by a similar adaptive regulation of the excitability, allowing a constrained reversible hypersensitivity state. A disruption of this adaptive activity regulation may lead to an unrestrained increase in PNs’ output, thus facilitating a transition from acute to pathological chronic pain.

## Results

### Medullary dorsal horn PNs reduce their excitability when acute pain is at its peak

Projection neurons (PNs) of the spinal and medullary dorsal horn integrate and summate noxious stimuli-evoked activity of peripheral neurons and local interneurons and relay its activity to higher brain centers^5^. Their firing, which determines the output from the spinal and medullary dorsal horn, depends on the synaptic input they receive and is refined by their intrinsic properties. Consequently, changes in the intrinsic properties of PNs, by affecting their output, could lead to alterations along the pain pathway, resulting in pathological pain. We, therefore, examined whether and how the intrinsic excitable properties of PNs change in acute and pathological chronic pain conditions. We identified medullary dorsal horn PNs by injecting mice with AAV_retrograde_-CamKII-tdTomato into the lateral parabrachial nucleus (PBN)^30^ (**Fig. 1a**). Targeted patch clamp recordings from labeled PNs (*see Methods*, **Fig. 1b)** revealed that the majority (70 out of 88 neurons, 80%) of the medullary dorsal horn PNs exhibited a delayed firing pattern (**Fig. 1c**).

**Fig 1.**
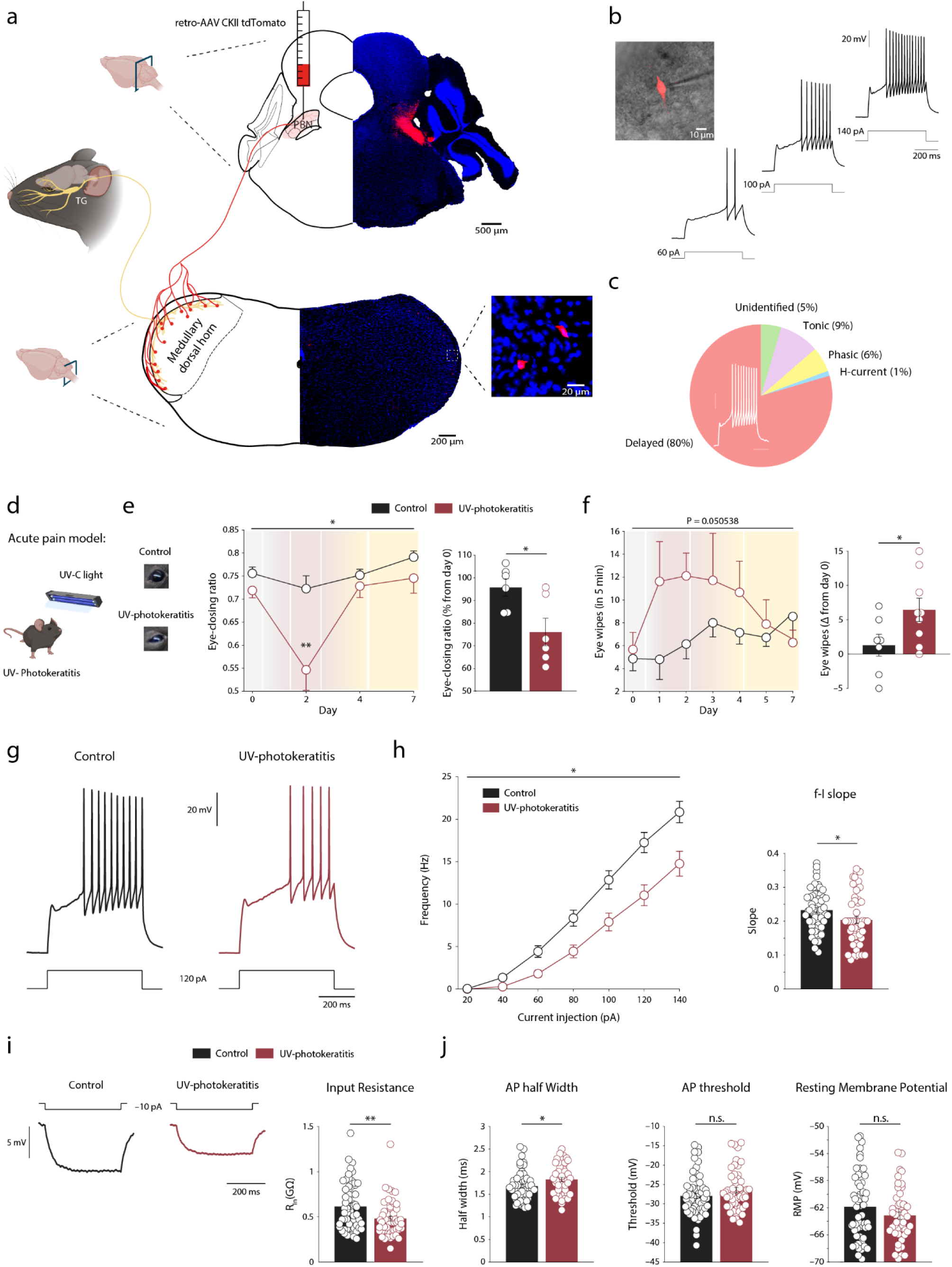
Medullary dorsal horn projection neurons decrease their excitability in acute pain. **a,** Schematic illustration and representative images of experimental procedure for retrograde labeling of medullary dorsal horn projection neurons (PNs). The retro-AAV CAMKII-tdTomato virus was injected into the parabrachial nucleus (PBN), and approximately three weeks after injections, labeled PNs were identified at the ipsilateral medullary dorsal horn of the trigeminal nucleus (TN). **b,** Representative image of tdTomato expressing PN with the recording pipette in acute medullary dorsal horn slice and representative traces of typical delayed firing pattern of the PN (representative of 70 out of 88). **c,** Distribution of medullary dorsal horn PNs based on their firing patterns, n cells/N mice: 88/29. Note that the majority of PNs possess delayed firing properties. **d,** Scheme depicting the induction of acute pain model of UV-photokeratitis (see *Methods*). **e,** Changes in eye-closing ratio during UV-photokeratitis-induced acute pain. *Left,* Representative images of the eye of mice in control and acute pain conditions two days after UV irradiation (*see Methods*). *Middle,* Eye-closing ratio at baseline (day 0), 2, 4, and 7 days after irradiation (*acute pain model, red*) or control procedure (*black*, see *Methods*, N=6 mice per group, *P_group_ =* 0.010879; *P_group×days_* = 0.037225). *Right*, Bar graphs of means ± s.e.m and individual values of the relative change (day 2 relative to day 0) in eye-closing ratio in acute pain and control conditions (N=6 mice per group, *P* = 0.0203). **f,** *Left,* Changes in the number of eye wipes following capsaicin application to the eyes of control (*black)* and acute pain (*red*) mice. Control: N = 7, UV-photokeratitis: N = 9*, P =* 0.050538. *Right,* Bar graphs of means ± s.e.m and individual values of the change in number of eye wipes between day 0 and day 2 (Control: N = 7, UV-photokeratitis: N = 9, *P* = 0.0484). **g,** Representative traces of action potential (AP) firing in response to a depolarizing current step recorded from identified PNs in slices from mice in acute pain (2 days after UV-irradiation, *red*) and control conditions (*black)*. Note decreased AP firing in PNs in acute pain conditions. **h,** *Left,* frequency-intensity (*f*-I) curves of PNs’ AP firing in acute pain and control conditions. n cells/N mice, Control: 60/26, UV-photokeratitis: 51/19, *P* = 0.014744. *Right,* Bar graphs of means ± s.e.m and individual values comparing the calculated *f*-I slope of PNs from both groups. Control: 61/26, UV-photokeratitis: 51/20, *P* = 0.0264. **i,** *Left,* Representative traces of changes in membrane voltage following hyperpolarizing current steps in PNs in control (*black)* and acute pain conditions (*red)*. *Right,* Bar graphs of means ± s.e.m and individual values comparing the calculated input resistance of PNs from both groups. n cells/N mice, Control: 59/26, UV-photokeratitis: 54/20, *P* = 0.0024. **j,** Comparison of the intrinsic excitable properties (specified above each panel; means ± s.e.m and individual values) of PNs from control (*black)* and acute pain conditions (*red).* n cells/N mice, *AP half-width*, Control: 62/26, UV-photokeratitis: 55/20, *P* = 0.012; *AP threshold*, Control: 62/26, UV- photokeratitis: 55/20, *P* = 0.1377; *RMP*, Control: 56/26, UV-photokeratitis: 49/20, *P* = 0.1374. Significance was assessed by two-way ANOVA for repeated measures with Tukey’s multiple comparisons test in **e** *(Middle)*; two-tailed unpaired student’s *t*-test in **e (***Right)*, **f** *(Right)*, **h** and **i**; linear mixed effects model (*see Methods)* in **f** *(Left) and* **g**. Data are presented as mean ± s.e.m. * *P* < 0.05, ** *P* < 0.01. n.s. not significant. Please refer to Supplementary Table 1 for full statistical information.

We then examined how the excitability properties of medullary PNs change in acute and chronic pain models. Here, we defined acute pain as a short-lasting and reversible pain. Therefore, we simulated acute pain using a UV corneal photokeratitis model^31, 32^ (**Fig. 1d**). Following UV irradiation, animals exhibited both ongoing pain (decreased eye-closing ratio of the eye) and hyperalgesia (increase in the number of eye wipes following capsaicin application), which peaked 2 days after irradiation and resolved after four days (**Fig. 1e,f**). Application of capsaicin to the eye two days after irradiation also led to an increased number of c-Fos expressing trigeminal ganglion (TG) neurons (**Supplementary Fig. 2**), suggesting that UV corneal photokeratitis leads to enhanced activation of primary sensory neurons innervating orofacial and craniofacial areas and conveying the information to the medullary dorsal horn.

Noxious stimuli-induced wiping is a non-reflexive complex response that requires activation of higher brain centers. The enhancement of this response and the appearance of ongoing pain strongly suggest that UV photokeratitis leads to amplified output from the medullary dorsal horn, resulting from the increased firing of PNs. However, contrary to our expectations, when acute pain and hyperalgesia peaked, PNs exhibited reduced firing and decreased gain (a sensitivity of a neuron to changes in input, **Fig. 1g-h**). This decrease in the firing of PNs was accompanied by decreased input resistance and increased AP half-width (**Fig. 1i-j**). No changes in AP amplitude, threshold, after-hyperpolarization, dV/dt_max_, resting membrane potential, or membrane capacitance were observed (**Fig. 1j** and **Supplementary Fig. 3)**. These results demonstrate that acute inflammation triggers a reduction in the excitability of PNs.

### Reduction in PNs’ excitability follows the timeline of acute pain

The decrease in the excitability of PNs could be attributed to the adaptive homeostatic regulation following increased activity, as was demonstrated in other systems^2–4^. In that case, the excitability of PNs should return to the baseline when pain hypersensitivity subsides. Indeed, on day 7 after UV irradiation, at which ongoing pain and hyperalgesia resolved (**Fig. 1e,f**), PNs did not show decreased firing, and PNs excitable parameters were similar to those of the control (**Fig. 2**). These data show that the excitable properties of PNs change along the time course of the acute pain, suggesting an adaptive regulation of excitability in medullary dorsal horn PNs.

**Fig. 2.**
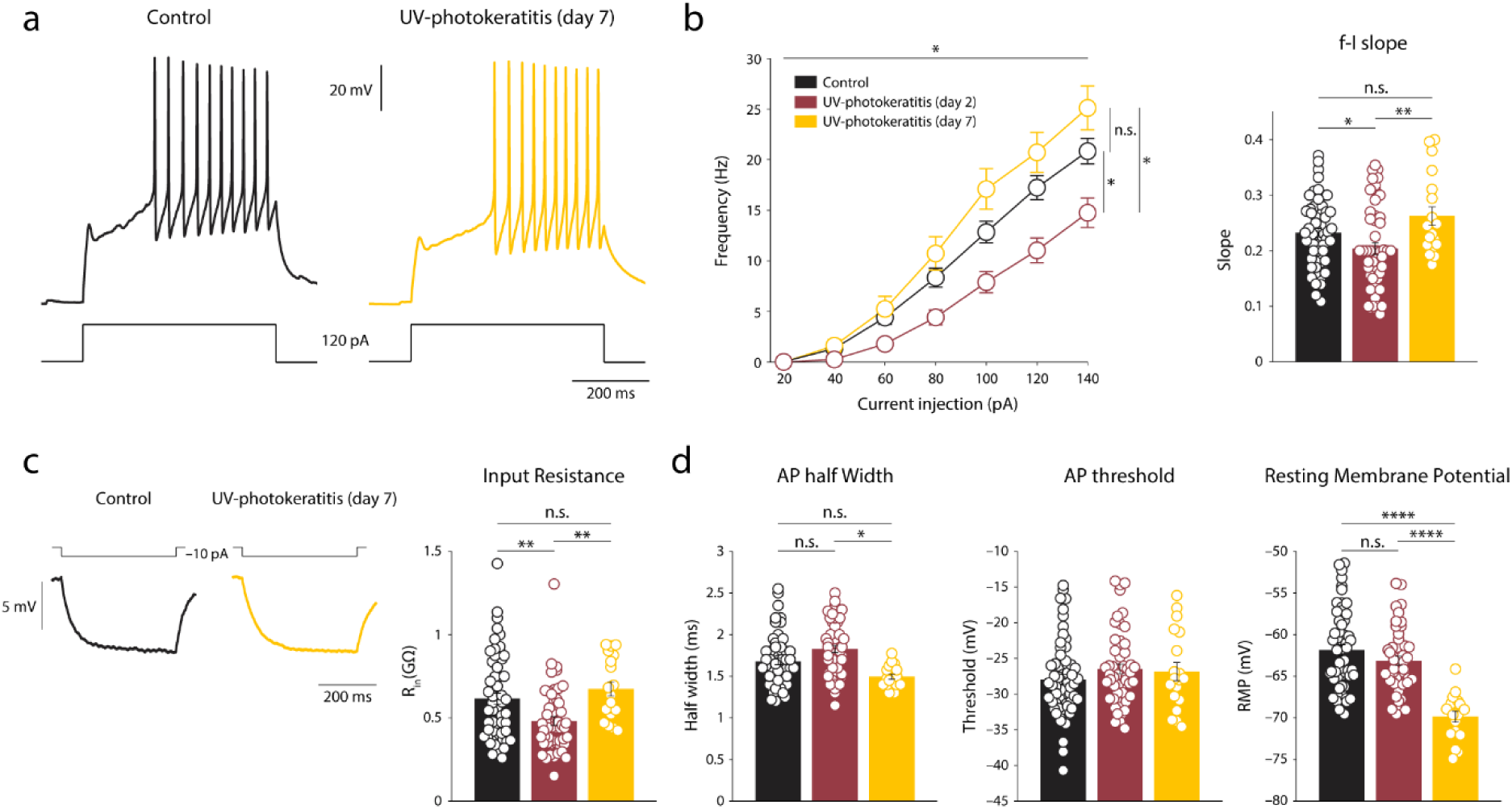
The change in PNs excitability follows the timeline of acute pain. **a,** Representative traces of AP firing in response to depolarizing current step of identified PNs in slices from control mice (*black)* and mice recovering from acute pain (*yellow,* 7 days following UV irradiation, see Fig. 1e, f). **b,** *Left,* Frequency-intensity (*f*-I) curves of PNs from the control (*black*), acute pain conditions (*red, data from* Fig. 1g), and recovery groups (7 days following UV irradiation, *yellow*). n cells/N mice, Control: 60/26, UV-photokeratitis: 51/19, UV-photokeratitis (day 7): 19/4, *P* = 0.029276. *Right*, Bar graphs of means ± s.e.m and individual values comparing the calculated *f*-I slope of PNs from all groups. Control: 61/26, UV-photokeratitis (day 2): 51/20, UV-photokeratitis (day 7): 19/4, *P* = 0.0097241. **c,** *Left,* Representative traces of PNs response to hyperpolarizing current steps in control mice (*black)* and mice recovering from acute pain 7 days following UV irradiation (*yellow)*. *Right,* Bar graphs of means ± s.e.m and individual values comparing the calculated input resistance of PNs from all groups. n cells/N mice, Control: 59/26, UV-photokeratitis (day 2): 54/20, UV-photokeratitis (day 7): 19/4, *P* = 0.0033831. **d,** Bar graphs of means ± s.e.m and individual values of intrinsic properties of PNs from control mice (*black)* and mice 2 days (acute pain, *red*) and 7 days (recovery, *yellow*) after UV irradiation. n cells/N mice*, AP half-width*, Control: 62/26, UV-photokeratitis (day 2): 55/20, UV-photokeratitis (day 7): 19/4, *P* = 0.049391; *AP threshold*, Control: 62/26, UV-photokeratitis (day 2): 55/20, UV-photokeratitis (day 7): 19/4, *P* = 0.35523; *RMP*, Control: 56/26, UV-photokeratitis (day 2): 49/20, UV-photokeratitis (day 7): 18/4, *P* = 2.4391e^-05^). In all panels, significance was assessed by fitting a linear mixed effects model (*see Methods)*. Data are presented as mean ± s.e.m.* *P* < 0.05, ** *P* < 0.01, **** *P* < 0.0001, n.s. not significant. Please refer to Supplementary Table 2 for full statistical information.

### The excitability of PNs is enhanced in chronic pain

We next asked whether a similar adaptive decrease in the excitability of PNs also occurs during pain hypersensitivity in chronic pain conditions. We examined the excitable properties of PNs in a model of chronic neuropathic pain, 2 - 4 weeks after inducing a chronic constriction to the distal infraorbital branch of the trigeminal nerve (CCI-dION) when mice exhibited substantial ongoing pain and mechanical allodynia (**Fig. 3b,c**) accompanied with increased activity of TG neurons^33–35^. Unlike acute pain conditions, PNs of CCI-dION mice significantly increase their intrinsic excitability (**Fig. 3d-g**). Notably, the changes in the excitability of PNs were precisely in the same parameters but in the opposite direction from the changes in acute pain, e.g., an increase in AP firing and neuronal gain, a decrease in AP half-width, and an increase in membrane input resistance (**Fig. 3d-g**). The increase in PNs’ excitability did not affect AP threshold, amplitude, after-hyperpolarization, dV/dt_max_, or resting membrane potential (**Fig. 3g** and **Supplementary Fig. 5**).

**Fig. 3.**
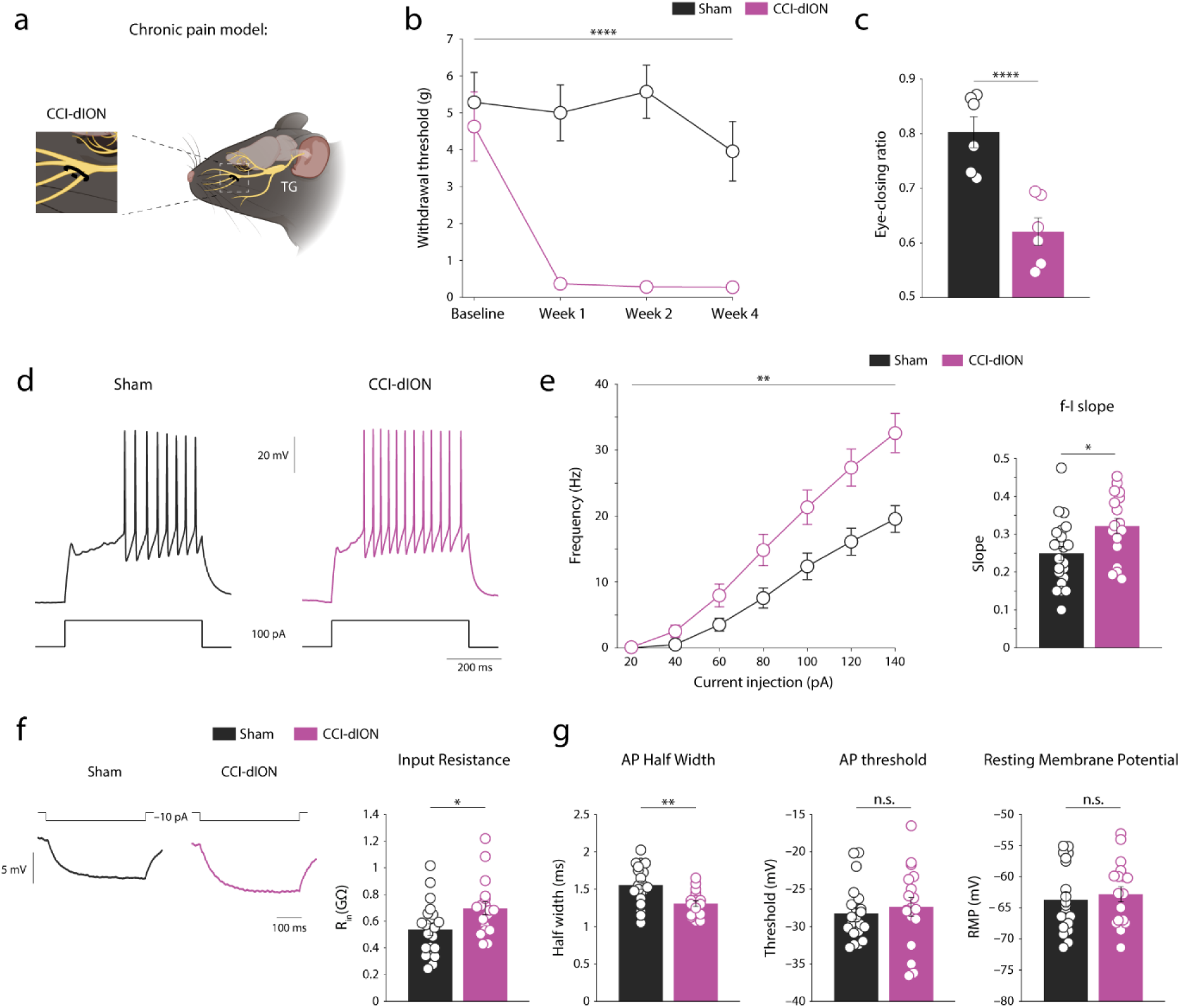
PNs increase their excitability in chronic neuropathic pain conditions. **a,** A scheme depicting the model used to establish chronic pain conditions (a chronic constriction injury of the distal infraorbital nerve, CCI-dION, *see Methods*). **b,** Changes in the sensitivity of sham (*black*) or CCI-dION mice (*purple*) to the application of mechanical stimuli measured as a withdrawal threshold to von Frey filaments applied to the mouse whisker pad (N=7 mice per group, *P_group_ =* 2.2214e^-06^; *P_group×time_* = 1.6972e^-07^). **c,** Eye-closing ratio of sham (*black)* or CCI-dION mice (*purple)* 4 weeks after surgery (N=6 mice per group, *P* = 7.382e^-04^). **d,** Representative traces of membrane voltage responses to depolarizing current step recorded from identified PNs in slices from mice in control (*sham*, 2-4 weeks after sham surgery, *black*) and chronic pain conditions (2-4 weeks after CCI-dION surgery, *purple*). Note an increase in action potential (AP) firing in chronic pain conditions. **e,** *Left,* Frequency-intensity (*f*-I) curves of PNs in chronic pain conditions and control mice. n cells/N mice, Sham: 22/4, CCI-dION: 18/4, *P =* 0.0031561. *Right,* Bar graphs of means ± s.e.m and individual values comparing the calculated *f*-I slope of PNs from both groups. Sham: 22/4, CCI-dION 18/4, *P* = 0.0128. **f,** *Left,* Representative traces of changes in membrane voltage following hyperpolarizing current steps in PNs in control (*black)* and chronic pain conditions (*purple)*. *Right,* Bar graphs of means ± s.e.m and individual values comparing the calculated input resistance of PNs from both groups. n cells/N mice, Sham: 21/4, CCI-dION: 18/4, *P* = 0.0204. **g,** Comparison of the intrinsic excitable properties (specified above each panel; means ± s.e.m and individual values) of PNs from control (*black)* and chronic pain conditions (*purple).* n cells/N mice, *AP half-width*, Sham: 22/4, CCI-dION: 18/4, *P* = 0.0013; *AP threshold*, Sham: 22/4, CCI-dION: 18/4, *P* = 0.5358; *RMP*, Sham 21/4, CCI-dION 18/4, *P* = 0.6035. Significance was assessed by two-way ANOVA for repeated measures with Tukey’s multiple comparisons test in **b**; two-tailed unpaired student’s *t*-test in **c**, **e,** and **f**; linear mixed effects model (*see Methods*) in **d**. Data are presented as mean ± s.e.m. * *P* < 0.05, ** *P* < 0.01, **** *P* < 0.0001, n.s. not significant. Please refer to Supplementary Table 3 for full statistical information.

Altogether, we show that the excitability of PNs undergoes adaptive regulation in acute but not chronic pain conditions, indicating differential modulation of the excitable properties of PNs in different pain states.

### Latency to the first AP determines the changes in PNs’ excitability in acute but not chronic pain

To screen for possible mechanisms underlying differential changes in PNs’ excitability, we further analyzed the electrophysiological data from the acute and chronic pain models. We showed that PNs decrease their firing rate in acute pain conditions and increase it in chronic pain conditions. Changes in the firing rate could involve modulations of various ion conductances and membrane properties. In our recordings, the calculation of the firing rate could be affected by changes in the latency to the first AP because AP frequency is calculated by dividing the AP number by the duration of the stimulation step. Longer latency will leave less time for APs to appear, producing a lower firing rate. Subsequently, the observed changes in firing may be explained by the changes in the biophysical mechanisms affecting the latency and not the firing properties. Notably, the vast majority of the medullary dorsal horn PNs display a prominent latency to the first AP (**Fig. 1b, c**), which persists at various stimulation intensities (**Fig. 1b**). We show in both models a strong negative correlation between the latency to the first AP and the firing rate (**Fig. 4a, d**). Therefore, we first examined whether the latency to the first AP changes in different conditions. In the acute pain model, the latency to the first spike in PNs was significantly higher compared to the control (**Fig. 4b**), suggesting that this change in latency could underlie the decreased firing rate. Indeed, when we calculated the firing rate just for the period of firing (see *Inset* and Methods), the difference in the firing between control and acute pain conditions was annulled (**Fig. 4c**). These data indicate that the decrease in the excitability of PNs in these conditions mostly depends on mechanisms governing AP latency.

**Fig. 4.**
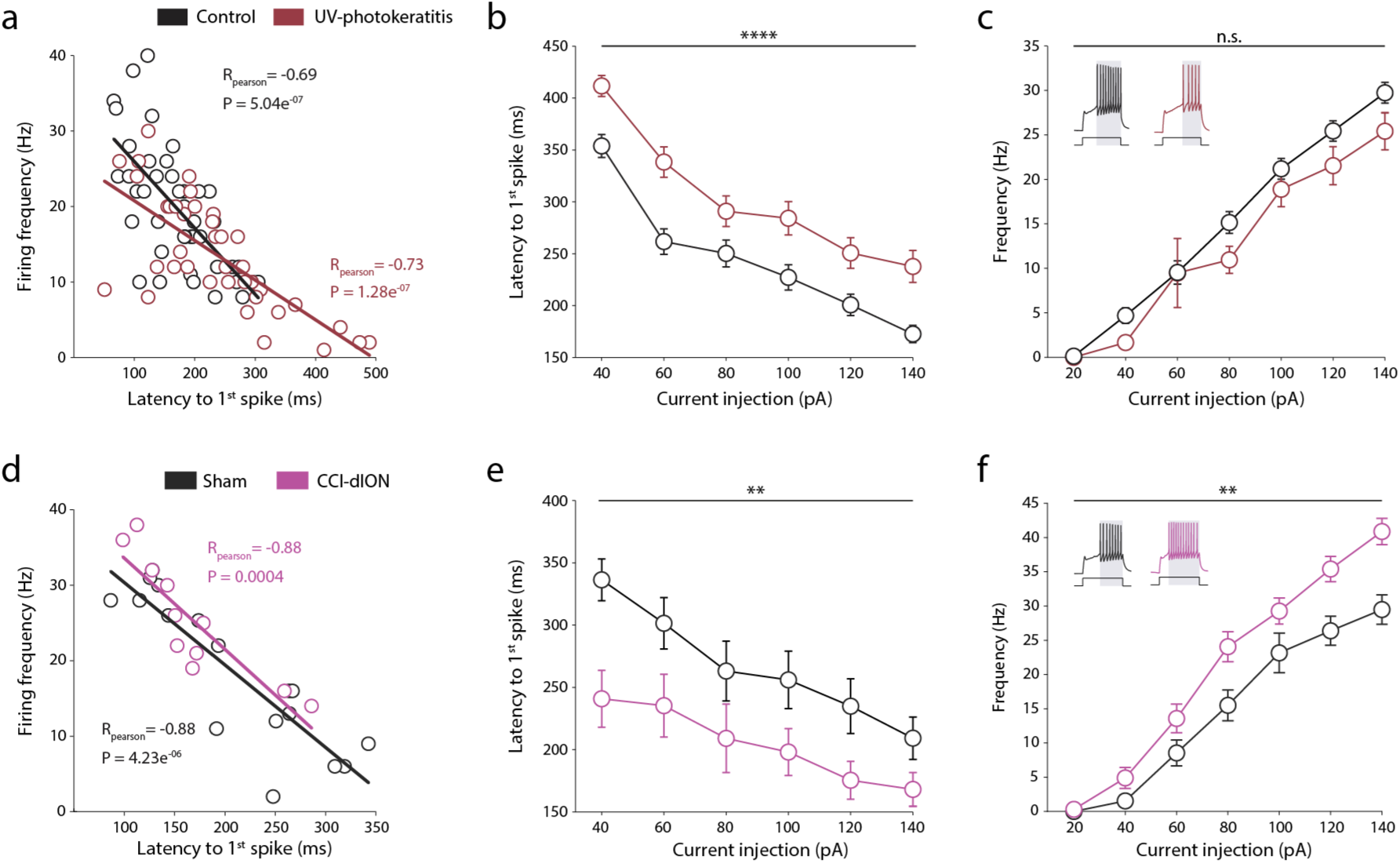
Changes in the latency to the first action potential affect PNs firing in acute but not chronic pain conditions. **a,** Correlations between the latency to the first action potential (first spike, measured at 140 pA current pulse) and the firing frequency of PNs in mice in acute pain (2 days after inducing UV photokeratitis, *red*) and control conditions (*black)*. Solid lines represent the linear regression of the data points. n cells/N mice, Control: 42/22, UV-photokeratitis: 38/19. **b,** Comparison of latencies to the first action potential (along the stimulation steps) in mice in acute pain (*red)* and control conditions (*black).* n cells/N mice, Control: 60/26, UV-photokeratitis : 47/19, *P =* 0.00023576. **c,** Frequency-intensity (*f*-I) curves in acute pain (*red)* and control conditions (*black)* calculated taking into account the changes in the latencies to the first action potential (by dividing the number of APs by the time from 1^st^ AP to the end of the pulse). Note that with this analysis, no difference in firing properties of PNs between acute pain and control conditions is observed (compare to Fig. 1g, *left*). n cells/N mice, Control: 60/26, UV-photokeratitis: 51/19, *P* = 0.087352. **d-f,** same as *a-c*, but the correlation and comparisons are between mice in chronic pain conditions (2-4 weeks after CCI-dION surgery, *purple*) and control mice (2-4 weeks after sham surgery, *black*). n cells/N mice**, d**, Sham: 17/4, CCI-dION: 11/3. **e,** Sham: 22/4, CCI-dION: 18/4, *P =* 0.0068754. **f,** Sham {: 22/4, CCI-dION: 18/4, group: *P* = 0.0029899. Significance was assessed by fitting a linear mixed effects model (*see Methods)* in **b, c, e** and **f**. Data are presented as mean ± s.e.m. ** *P* < 0.01, **** *P* < 0.0001, n.s. not significant. Please refer to Supplementary Table 4 for full statistical information.

In the chronic pain conditions, the latency for the first AP was significantly shorter than in the sham group (**Fig. 4e**). Notably, the increase in AP firing remained after correcting for the AP latency (**Fig. 4f**), suggesting that other mechanisms beyond changes in the latency underlie the increase in PNs excitability in the chronic pain model.

Altogether, these data suggest that the decrease in the excitability of PNs in acute pain conditions depends, at least in part, on the mechanisms governing the latency for the first spike. The increase in PNs excitability in chronic pain is likely to be mediated by different mechanisms.

### Potassium A-current I_A_ increases in acute but not chronic pain conditions

It has been previously demonstrated that delayed firing is modulated by the activity of Kv4 isoforms of voltage-gated potassium channels underlying I_A_-current (I_A_)^36^. Here, we show that in medullary dorsal PNs, the blockade of I_A_ by 4-aminopyridine (4-AP)^37^ significantly reduced the latency to the first spike (**Fig. 5a**). Because we showed that the increase in latency was the main factor leading to decreased PNs’ firing, we hypothesized that the increase in I_A_ could underlie the decrease in the excitability of PNs in acute pain conditions. We isolated and analyzed I_A_ in PNs and examined whether and how it changed when acute pain and hyperalgesia peaked, i.e., 2 days after corneal UV irradiation (**Fig. 5b**, see *Methods*). We show that although the voltage dependence of I_A_ activation did not change (**Fig. 5b and Supplementary Fig. 7c**), the voltage dependence of inactivation was right-shifted (**Fig. 5b, c)**, leading to an increased “window” current (**Fig. 5d**). Because the “window” current defines the availability of the channels, our data indicate that in acute pain conditions, the Kv4 channel availability around the resting membrane potential increases. Furthermore, we show that in acute pain conditions, the fast component of the I_A_ decay time became substantially slower (**Fig. 5e, f**), implying an increase in overall current. Indeed, we show that in acute pain conditions, I_A_ charge (*see Methods*) was significantly higher (**Fig. 5g**).

**Fig. 5.**
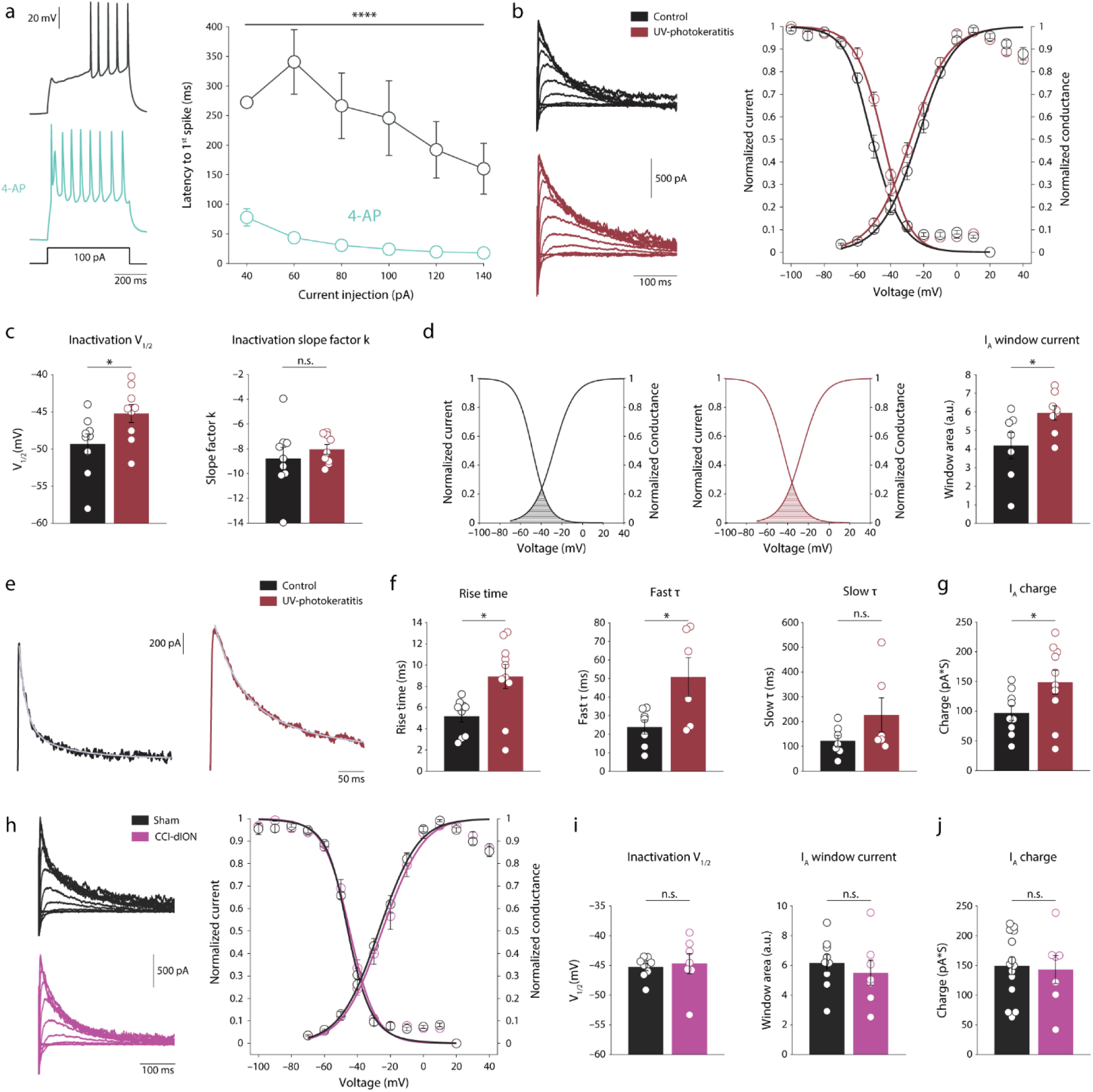
A-type potassium current in PNs increases in acute but not chronic pain conditions. **a,** *Left,* Representative traces of AP firing responses of the same PN before (*above*) and after (*below*) application of 5 mM 4-aminopyridine (4-AP). *Right,* Comparison of latencies to first action potential in the same PNs before and after application of 4-AP (n cells/N mice: 6/2, *P =* 3.4976e^-17^). **b,** *Left,* Representative traces of the isolated I_A_ in PNs in slices from mice in acute pain (*red)* and control conditions (*black).* For the details of I_A_ isolation, please refer to the Methods and Supplementary Fig. 6. *Right,* Voltage dependence of I_A_’s activation and inactivation in acute pain (*red)* and control conditions (*black)*. Data were fit by Boltzmann functions (see Methods). n cells/N mice, *Activation curves*, Control: 9/7, UV-photokeratitis: 10/4; *Inactivation curves*, Control: 9/7, UV-photokeratitis: 9/4. **c,** V_1/2_ (*left*), and slope factor k (*right)* of the I_A_ inactivation curve in acute pain (*red)* and control conditions (*black).* n cells/N mice, Control: 9/7, UV-photokeratitis: 9/4, V_1/2_: *P* = 0.0405; slope factor k: *P* = 0.4636. **d,** *Left,* Representative activation and inactivation fitted curves of individual PNs in acute pain (*red)* and control conditions (*black)*. The shaded area under the curves is defined as the window current. *Right,* I_A_ window current of PNs in acute pain (*red)* and control conditions (*black).* n cells/N mice, Control: 7/7, UV-photokeratitis: 8/4. *P* = 0.041. **e,** Representative traces of the isolated I_A_ in response to a voltage step to +10 mV in acute pain (*red)* and control conditions (*black)*. Superimposed gray traces are the 2-exponential fit to the decay of the current. **f,** Comparison of I_A_ kinetics (indicated above the panels) in acute pain (*red)* and control conditions (*black).* n cells/N mice*, Rise time*, control: 9/7, UV-photokeratitis: 10/4. *P* = 0.0113; *Fast τ*, control: 7/7, UV-photokeratitis: 6/4. *P* = 0.0492; *Slow τ*, control: 7/7, UV-photokeratitis: 6/4. *P* = 0.1981. **g,** Comparison of the I_A_ current charge at stimulation to +10 mV in acute pain (*red)* and control conditions (*black).* n cells/N mice, Control: 9/7, UV-photokeratitis: 10/4. *P* = 0.0485. **h-j,** same as *b, c,* and *g* but compared in PNs in chronic pain and sham conditions. n cells/N mice, *Activation curves*, sham: 14/5, CCI-dION: 7/5; *Inactivation curves*, Sham: 9/5, CCI-dION: 7/5; *I_A_ properties*: Sham: 9/5, CCI-dION: 7/5, *Inactivation V_1/2_*: *P* = 0.7468; window current, *P* = 0.5185; *I_A_ charge*: Sham: 14/5, CCI-dION: 7/5. *P* = 0.8171. Significance was assessed by fitting a linear mixed effects model (*see Methods)* in **a** *right*; two-tailed unpaired student’s *t*-test for **c, d, g, i** *(Right)* and **j**; two-tailed unpaired student’s *t*-test with unequal variance for **f** and **i** (*Left)*. Data are presented as mean ± s.e.m. * *P* < 0.05, **** *P* < 0.0001, n.s. not significant. Please refer to Supplementary Table 5 for full statistical information.

Importantly, in the model of CCI-ION-induced chronic pain, I_A_ properties and kinetics did not change (**Fig. 5h-j** and **Supplementary Fig. 8a-c**).

Our data indicate that I_A_ current increases in acute but not chronic pain conditions.

### Changes in I_A_ kinetics in acute pain conditions are sufficient to produce a decrease in the AP firing of PNs

We next examined whether the increase in I_A_ in acute pain conditions could underlie the reduction in PNs excitability. We built a point-neuron computational model of PN based on the previously described model of PN^38^. We added a module of I_A_, with properties taken from our experimental data, and fit the parameters to replicate the characteristic firing of PNs we described here (*see Methods*, **Fig. 6a**, see also **Fig. 1b**). We then simulated the shift of I_A_ voltage dependence of inactivation in a range of values of V_1/2_ (the membrane potential at which 50% of the channels are in the inactive state) measured at the control conditions and culminating in the values we measured in the acute pain conditions and examined the changes in the firing of the simulated PN (**Fig. 6**). We show that the rightward shift in the V_1/2_ of I_A_ inactivation curve produces a gradual decrease in AP firing frequency and an increase in the latency to the first AP (**Fig. 6c, e**). Notably, at the V_1/2_ values measured in acute pain conditions, the changes were similar to the phenomenon we observed in the experimental conditions (compare **Fig. 6a-b, d** to **Fig. 1g** and **Fig. 4b**). These results predict that the rightward shift in the I_A_ inactivation we measured in acute pain conditions is sufficient to produce the decrease in the excitability of PNs. Altogether, we show that the excitability of PNs undergoes I_A_-mediated adaptive regulation in acute pain conditions.

**Fig. 6.**
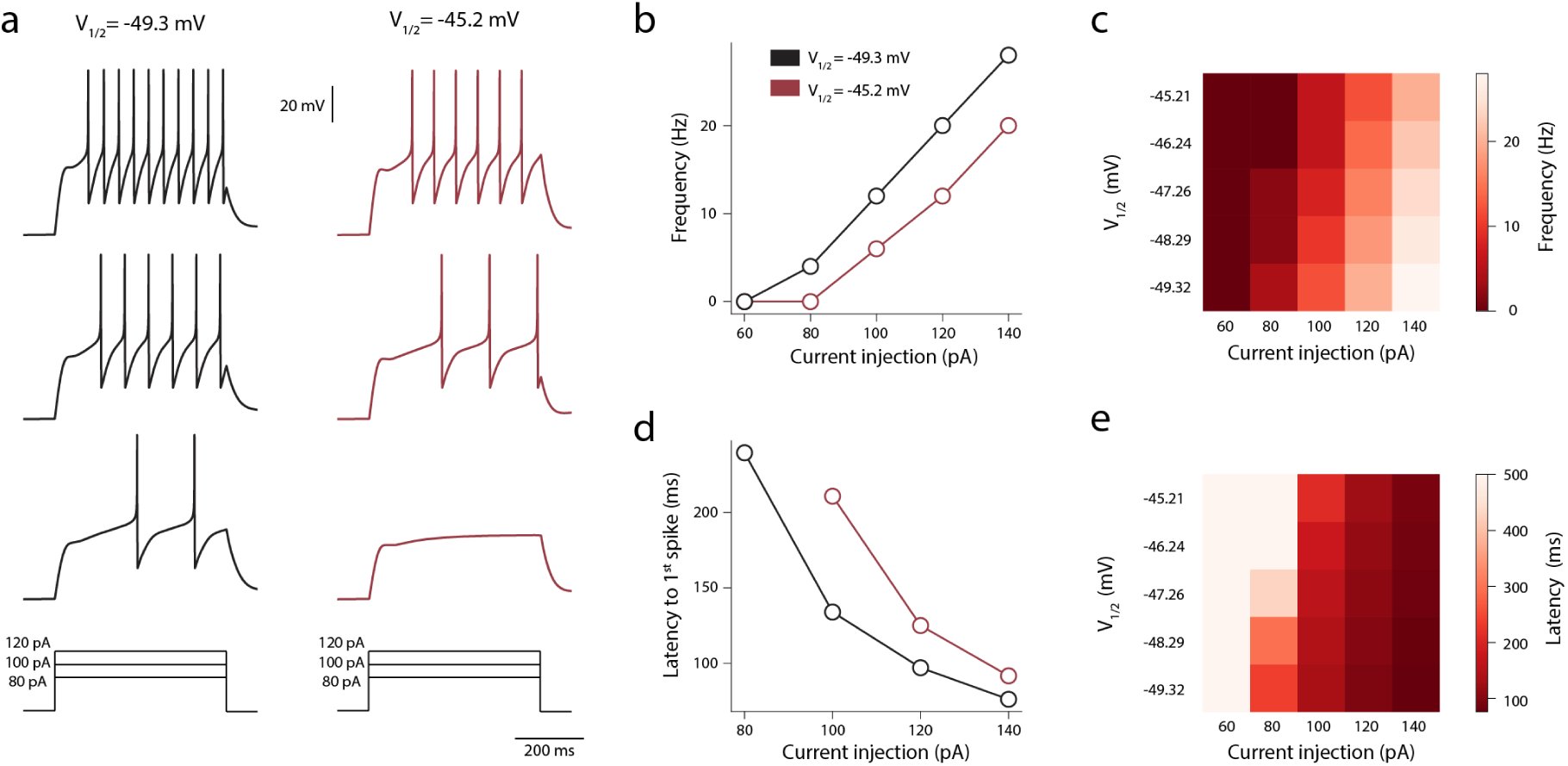
A rightward shift in the inactivation curve of I_A_ is sufficient to reduce AP firing. **a,** Representative traces of firing responses from a modeled PN in simulated acute pain (*red)* and control conditions (*black)*, i.e., when the V_1/2_ of the I_A_ inactivation was set to the values experimentally measured in acute pain (V_1/2_ = −45.2 mV) or control conditions (V_1/2_ = −49.3 mV, see Fig. 5c). **b,** Frequency-intensity (*f*-I) curves (calculated for the time of entire stimulation) from modeled PN in simulated acute pain (*red)* and control conditions (*black)*. **c,** The relation between voltage dependence of I_A_ inactivation and PNs firing at different stimulations. Note that the depolarizing shift of V_1/2_ of I_A_ inactivation leads to a decrease in PNs firing. **d,** Latency to the 1^st^ AP for different stimulation intensities from modeled PN in simulated acute pain (*red)* and control conditions (*black)*. **e,** The relation between voltage dependence of I_A_ inactivation and the latency to 1^st^ AP at different current intensities. Note that the depolarizing shift of V_1/2_ of I_A_ inactivation leads to an increase in the latency to 1^st^ AP.

### In acute pain conditions, the number of activated PNs increases

Our results demonstrate that in acute pain conditions, individual PNs decrease their firing. On the other hand, our behavioral results suggest increased overall output from the PNs toward higher brain centers. If the number of activated PNs increases in acute pain conditions, this could still result in increased output despite the reduction in excitability of individual cells. To examine this hypothesis, we assessed the number of activated PNs in acute pain conditions by measuring the number of PNs expressing c-Fos two days after inducing UV-photokeratitis. Because the changes in the excitability of PNs we described occur spontaneously, i.e., without applying any exogenous stimuli to the cornea, we first assessed the number of c-Fos expressing PNs without applying noxious stimuli. We show that the number of activated PNs was significantly higher in acute pain conditions than in control (**Fig. 7a, b**). After applying capsaicin to the cornea, the number of activated PNs was also significantly higher than in the control (**Fig. 7c, d**). The increased number of activated PNs could underlie the expected amplification of overall output from the dorsal horn despite the reduction in the excitability of individual PNs during inflammatory pain.

**Fig. 7.**
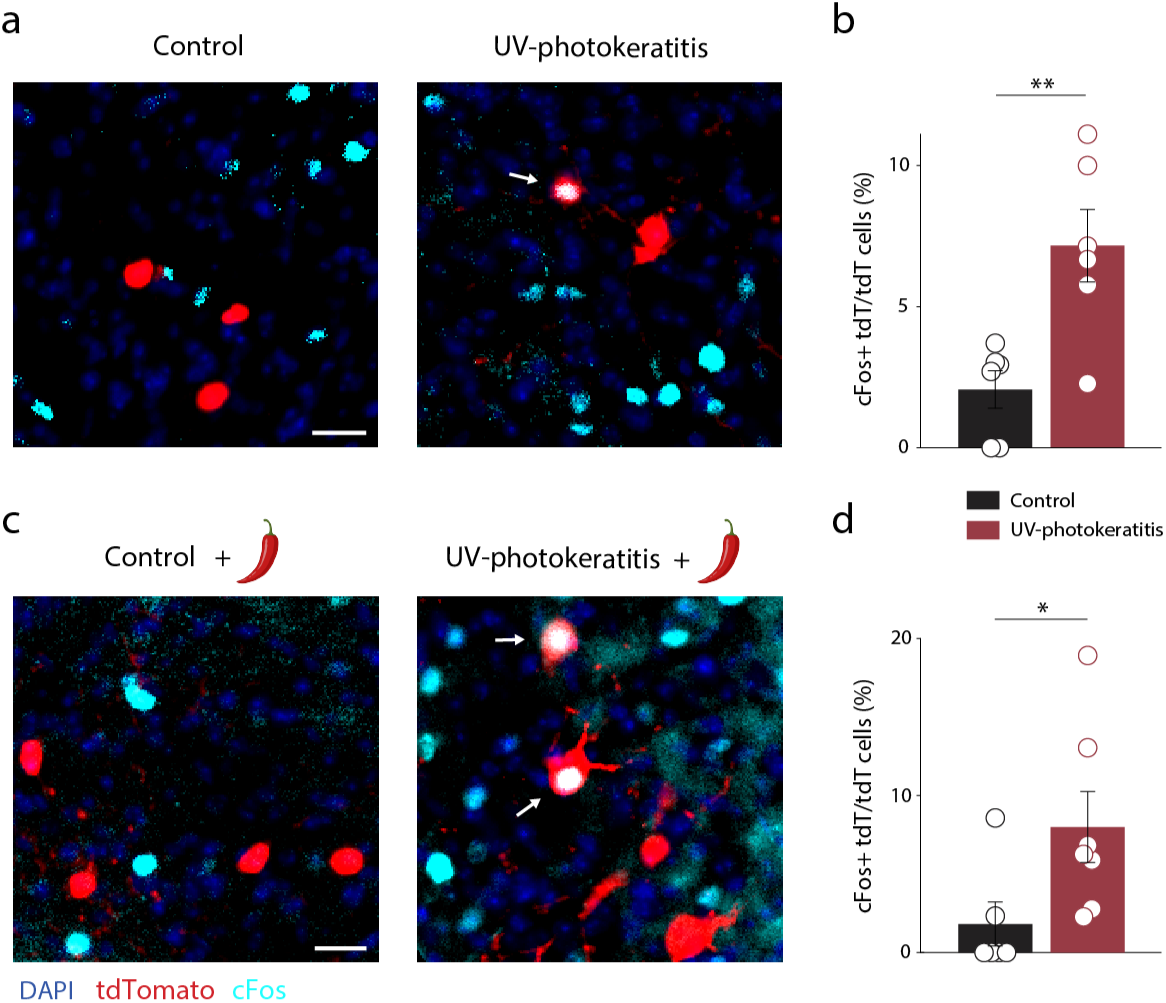
The number of activated PNs increases in acute pain conditions. **a,** Representative images of retrogradely labeled PNs stained for c-Fos in medullary dorsal horn slices from mice in acute pain and control conditions that did not undergo any stimulation. The arrow indicates a co-labeled (tdTomato and c-Fos) PN. Scale bar: 20 µm. **b,** Bar graph of means ± s.e.m and values from the individual mice of the percentage of activated PNs (expressing tdTomato and c-Fos) among all PNs (expressing only tdTomato, N = 6 mice per group, 4-5 slices per mouse; *P* = 0.0056). Each data point represents the mean percentage of co-labeled PNs measured from individual mice. **c, d,** same as *a, b* but from mice after corneal application of capsaicin. Control: N = 6, UV-photokeratitis: N = 7; 3-5 slices per mouse. *P* = 0.0475). Significance was assessed by a two-tailed unpaired student’s *t*-test in **b** and **d**. Data are presented as mean ± s.e.m. * *P* < 0.05, ** *P* < 0.01. Please refer to Supplementary Table 6 for full statistical information.

## Discussion

Regulatory homeostatic processes maintain neurons within a narrow activity range despite changing conditions. Studies in recent years suggest that chronic neurological disorders such as Alzheimer’s disease, epilepsy, and neuropathic pain should be viewed as failures in homeostatic regulation^39–41^. Accordingly, therapies for these diseases should be targeted to assist in regaining the original setpoint by searching for the processes preventing proper homeostatic regulation. Here, we show that homeostatic regulatory processes occur in medullary dorsal horn PNs during acute but not chronic pain. Although both types of injury lead to increased activity of the primary afferent ^31, 33–35^ (also see **Supplementary Fig. 2**), the excitability of medullary dorsal horn PNs reversibly decreases during acute inflammation but non-reversibly increases during chronic neuropathic nerve injury.

In theory, in normal conditions, the output of PNs is directly related to the input they receive from the primary afferents (**Supplementary Fig. 9**, *upper*). Consequently, when the input from the periphery increases in inflammatory conditions, the output from PNs should increase proportionally. However, our data suggest that the overall increase in output from PNs in acute inflammatory pain conditions (by factor “*a**’***,” **Supplementary Fig. 9**, *middle*) is not proportional but is tuned by the homeostatic decrease in PNs gain, i.e., decreased sensitivity of PNs to the changes in input (by factor “*b*,” **Supplementary Fig. 9**, *middle*). This would ultimately lead to a “down-tuning” of single PNs’ output (“*a**’**/b*,” **Supplementary Fig. 9**, *middle*), which would regulate the overall output, plausibly minimizing upstream plastic changes along the pain neuroaxis and preventing pain chronification. The lack of adaptive regulation and the increase in PNs excitability (factor “*d*,” **Supplementary Fig. 9**, *bottom*) together with increased peripheral input (factor “*c*,” **Supplementary Fig. 9**, *bottom*) may lead to “unrestrained” output (“*c**’**\*d*”) from the dorsal horn, plausibly leading to nociplastic changes and pain chronification (**Supplementary Fig. 9**, *bottom*).

The adaptive decrease in intrinsic excitability of PNs during the peak of inflammatory hyperalgesia and pain is counterintuitive as hyperalgesia and ongoing pain suggest an increased output to higher centers. The decrease in the gain of PNs suggests that in acute pain conditions, PNs will produce less firing for a specific input, but the resulting output could still be higher than the control (“*a**’**/b*”>1, **Supplementary Fig. 9**, *middle*). The increased number of activated PNs following inflammation (**Fig. 7c, d**) can also lead to increased output despite the decrease in excitability of the individual PNs. Homeostatic regulation of neuronal excitability is a well-studied phenomenon known to be initiated by increased synaptic or intrinsic neuronal activity^2–4^. Similarly, we showed an increased number of c-Fos expressing PNs during acute pain even without applying noxious stimuli (**Fig. 7a, b**), probably attributable to increased input from primary afferents^31, 32^ (see also **Supplementary Fig. 2)** and local interneurons^20, 42^. We posit that this increase in the activity of PNs could trigger the adaptive decrease in intrinsic excitability described here.

We used a rational analysis of the electrophysiological data to identify a potential biophysical correlate of the decrease in the excitability of PNs in acute pain. The lack of changes in AP amplitude, threshold, and maximal rate of change of AP voltage (dV/dt_max_) excluded the modulation of voltage-gated sodium channels, and the decrease in input resistance ruled out axon initial segment-mediated effects^3^. On the other hand, the prolongation of the latency to the first AP^36^ (see also **Fig. 4b and Fig. 5a**) and the increase in AP half-width^43^ indicated that modulation of the potassium I_A_ current might play a role in the inflammation-induced decrease in the excitability of PNs. Indeed, we show a rightward shift of the voltage dependence of I_A_ inactivation, increasing I_A_ availability and total charge, leading to longer delays. Moreover, our computational model predicts that the rightward shift in the voltage dependence of I_A_ inactivation decreases membrane input resistance by about 40 MΩ, plausibly by increasing the number of available Kv4 channels.

The role of I_A_ in defining the excitability of dorsal horn interneurons in normal and pathological conditions is well established^44, 45^. Recently, an increase in I_A_ following the release of TAFA4 cytokine from low threshold C-fibers into the dorsal horn was suggested as one of the mechanisms for the decrease in the excitability of inhibitory dorsal horn neurons in chronic pain conditions^45^. Further, substance P (SP)-induced modulation of I_A_ in striatal cholinergic neurons was found to affect their dendritic excitability^46^. Considering that PNs express the receptors for SP, NK1^5^, it is plausible that enhanced SP release from primary peptidergic nociceptors during acute pain underlies the I_A_ increase in PNs. Other possible candidates for modulating I_A_ in PNs are dynorphin, released from local interneuron, or oxytocin from the hypothalamus-to-spinal cord projections, both of which have been shown to upregulate I_A_ in dorsal horn neurons^47, 48^. Nevertheless, our results do not exclude that modulation of other conductances could contribute to the adaptive decrease in PNs’ excitability. Unlike the acute pain model, the changes in neuronal activity in chronic pain conditions were not associated with the downregulation of I_A_, and PNs increase their evoked firing and excitability. This may contribute to the increase in the output of PNs following chronic neuropathic injury^49^ and could potentially trigger plastic, sensitization-like changes leading to pain chronification.

Our data suggest that the neuropathic-induced changes in the excitability of PNs are driven by mechanisms different from those in the acute pain model. In both cases, the input resistance of PN changes: it decreases in acute pain conditions but increases in chronic pain conditions (**Figs. 1i and 3f**). The analysis of the correlation between the changes in input resistance and firing shows a minor correlation between these factors in acute pain conditions but a strong correlation in chronic pain conditions (**Supplementary Fig. 6**). One of the factors affecting input resistance is cell size. Changes in cell size would result in changes in membrane capacitance. This was not the case for PNs in acute pain conditions (**Supplementary Fig. 3**). On the other hand, in chronic pain conditions, we show a significant decrease in the membrane capacitance of PNs (**Supplementary Fig. 5**), suggesting a decrease in cell size which could explain the observed increase in the input resistance. The nerve injury-induced changes in the morphology of infralimbic layer 2/3 pyramidal cells, affecting their capacitance, were previously reported in chronic pain^50^. In addition, inhibition of potassium Kv7/M current^51^ or inward rectifying potassium leak conductance^52^ could explain both the increase in input resistance and PNs excitability observed in chronic pain model^53, 54^.

One limitation of our study is that all the data were collected from male mice. We started the experiments in both sexes, but surprisingly, female mice did not show corneal hypersensitivity following UV irradiation (**Supplementary Fig. 1**). We, therefore, continued our experiments on male mice.

In summary, our results reveal an adaptive mechanism, which, when absent, could produce pain chronification. In acute pain conditions, when the input from primary sensory neurons increases, PNs exhibit I_A_-mediated adaptive decrease in their intrinsic excitability, resulting in reduced PNs’ firing. The reduction in PNs’ firing balances the response of PNs to the increased input from the periphery, thus controlling overall output from the medullary dorsal horn towards higher brain centers. In chronic pain conditions, the activity of PNs is not balanced by an adaptive I_A_ modulation, and PNs show elevated intrinsic excitability. The elevated extrinsic inputs “falling” on membranes of PNs with increased intrinsic excitability could produce excessive firing, amplifying the output from the dorsal horn, thus triggering pain chronification.

## Methods

### Animals

Adult (9-13 weeks) C57BL/6J mice were used in all experiments unless stated otherwise. Animals were group-housed under a controlled temperature (21 ± 2°C) and humidity (55 ± 10%) on a 12h-12h light/dark cycle, with *ad libitum* access to food and water. All experimental procedures were approved by the Institutional Animal Care and Use Committees at the Hebrew University of Jerusalem.

Initially, we planned to perform all the experiments on both sexes. We started by examining the development of acute pain following inducing UV-photokeratitis. We examined the responses of both male and female mice to capsaicin application to the cornea following UV irradiation. Surprisingly, in our hands, female mice did not develop inflammatory hypersensitivity following UV irradiation (**Fig. 1f** and **Supplementary Fig. 1**). Therefore, we did not include female mice for the rest of the experiments in this study.

### Pain models

#### Acute pain model: UV-photokeratitis model

Corneal photokeratitis was induced as previously described^31, 32^. Briefly, mice were anesthetized with a combination of 70 mg/kg ketamine and 0.8 mg/kg medetomidine, intraperitoneally (i.p.). Mice were placed on a heating pad, and one eye was exposed to UV-C radiation (254 nm, 450 mJ/cm^2^), delivered through a UV light source (VL-4.C 230V 50/60Hz; Viber Lourmoat, Marne-la-Vallee, France), from a distance of 17 cm for 22 minutes. Following UV radiation, mice were injected with 1.3 mg/kg atipamezole for anesthesia reversal. Control mice underwent the same procedure but without the UV radiation.

#### Chronic pain model: Chronic constriction injury of the distal infraorbital nerve (CCI-dION)

The CCI-dION model was performed as previously described^55, 56^. In short, mice were anesthetized with a combination of 70 mg/kg ketamine and 0.8 mg/kg medetomidine (i.p.) and subcutaneous injection of 1-2 mg/kg Meloxicam for analgesia. Then, mice were placed on a heating pad, and the skin below the infraorbital foramen (3 mm) was shaved and cleansed using 10% betadine. A 0.5 cm incision was made, and the nerve was gently exposed by blunt preparation. A chromic gut ligature (5-0) was loosely tied around the distal part of the ION. The skin incision was sutured, and mice were injected with 1.3 mg/kg atipamezole for anesthesia reversal. Sham mice underwent the same procedure of nerve exposure but without ligation.

### Behavioral tests

All mice were handled for 5 consecutive days before the behavioral experiments. For 3 days before the experiments, as well as during experimental days, mice were habituated to head restriction by the experimenter and to the behavioral room (≥30 min) and recording chambers used (≥10 min). The experimenter was blinded to mice groups in all the experiments and analyses.

#### Assessment of ongoing pain: Eye-closing ratio

The eye-closing ratio is a measurement of the orbital tightening used to quantify ongoing pain^57–59^. The eye-closing ratio was calculated as the ratio of the height and width of the tested eye. The height was defined as the distance between the edge of the upper and lower eyelids, and the width as the distance between the internal and external canthus. For eye-closing ratio measurements over the course of corneal UV-photokeratitis, in all experimental days (0, 2, 4, and 7), mice were head-restricted and photographed. For eye-closing ratio measurements in the CCI-dION model, measurements were done 4 weeks following the CCI-dION / sham procedure.

#### Assessment of nociceptive hyperalgesia

To assess the changes in sensitivity to a noxious stimulus, behavioral responses produced by corneal instillation of capsaicin were recorded under control and acute pain conditions. After habituation, mice were treated with a 10 µl drop of the vehicle onto the tested eye (0.1% ethanol in standard extracellular solution, SES). SES was composed of 145 mM NaCl, 5 mM KCl, 2 mM CaCl_2_, 1 mM MgCl_2_, 10 mM glucose, and 10 mM HEPES) and video recorded for 5 minutes in the recording chamber (a custom-made 10×20×25 cm transparent plexiglass). 1 hour later, 10 µl of 1 µM capsaicin (in SES) was applied to the tested eye, and the mouse was video recorded for 5 minutes in the recording chamber. Eye wipes were manually counted post hoc. Mice (N=2) that showed apathetic behavior manifested by reduced motor activity were noted by the blinded experimenter and were excluded from the analysis. Mice (N=2) that showed no eye wipe responses to both vehicle and capsaicin in baseline (day 0) were excluded from the analysis. In the trials in which mice were not in front of the camera, preventing correct detection of eye wipes for at least 30 seconds out of the first minute following capsaicin application were excluded from analysis.

#### Assessment of mechanical allodynia

To assess changes in sensitivity to innocuous mechanical stimulus, withdrawal responses to von Frey filaments applied to the middle area of the whisker pad were measured in mice in sham and CCI-dION conditions. Mechanical sensory thresholds were assessed using von Frey filaments of 0.07g to 6g by the simplified up-down (SUDO) method while immobilizing the mice through neck pinch^60^. A positive response was recorded if the mouse withdrew its head in 3 out of 5 trials for each filament.

### Stereotactic injections

5-6 weeks old mice were anesthetized with a combination of 60 mg/kg ketamine, 0.5 mg/kg medetomidine, and 0.4-0.6% isoflurane (i.p.), pretreated with a subcutaneous injection of 1-2 mg/kg Meloxicam and placed on a heating pad in a stereotactic frame (KOPF instruments). The cranium was shaved, and the skin was cleansed using 70% ethanol and 10% betadine. A 1-1.5 cm midline incision was made, followed by drilling a small hole in the scalp using a fine drill burr. Unilateral injections to the PBN were delivered at the following coordinates (from bregma): ML: 1.3 mm, AP: −5.2 mm, DV: −3.75 mm using a microliter syringe (35g nanofil needle, World Precision Instruments, Sarasota, FL) loaded with retro-AAV-CaMKII-tdTomato (titer 4.5×10^12^ genome copies per ml, ELSC vector Core Facility, Israel). 250 nl of the virus was injected at 100nl/min via UltraMicroPump (World Precision Instruments, Sarasota, FL). The needle was left in the tissue for an additional 5-10 minutes before being slowly retracted. Finally, the incision was sutured using chromic gut ligature (5-0) or glued by tissue adhesive (VetBond), and 1 mg/kg atipamezole was injected for anesthesia reversal.

### Electrophysiological recordings

#### Slice preparation

3-7 weeks after injections of the retro-AAV-CaMKII-tdTomato, mice were deeply anesthetized with isoflurane, decapitated and the brain stem was quickly removed into a warm (34°C) artificial cerebrospinal fluid (aCSF) solution containing (in mM): 126 NaCl, 3 KCl, 1.3 MgSO_4_, 1.2 KH_2_PO_4_, 26 NaHCO_3_, 10, 2.4 CaCl_2_, saturated with 95% O_2_/5% CO_2_. 300 µm coronal slices of caudal brain stem containing the medullary dorsal horn were prepared using a vibratome (VT 1200S, Leica). Slices were incubated in warm (34°C) aCSF solution for 1 hour, after which they were maintained at room temperature (23 ± 2°C) until recording.

#### Whole-cell recordings

Slices were transferred into a recording chamber and continuously perfused with oxygenated aCSF (the same as cutting solution). PNs of the medullary dorsal horn were visualized using differential interference contrast infrared microscopy (BX61WIF, Olympus, Tokyo, Japan) and identified by location and tdTomato expression using a CoolLed fluorescence excitation system. Borosilicate fire-polished patch pipettes (4-6 MΩ, pulled on a P-1000 puller, Sutter, USA) were used for all experiments. Signals were amplified using Multiclamp 700B (Molecular Devices) and digitized at 20-50 kHz (and low-pass filtered at 3 kHz) with Digidata 1440A (Molecular Devices) interfaced with pClamp 10.3 (Molecular Devices). Electrode series resistance (R_s_) was routinely checked, and recordings were discarded if: (1) R_s_ was more than 30 MΩ; (2) R_s_ changed >15% during recordings, and (3) if resting membrane potential (RMP) was higher than −50 mV.

#### Current clamp recordings

The intrinsic properties of PNs were recorded using internal solution containing either (in mM): 130 K-gluconate, 4 Na_2_ATP, 0.5 NaGTP, 20 HEPES, 0.5 EGTA, 5 KCl (internal solution 1, 283 mOsm, pH was adjusted to 7.3 with KOH), or (in mM): 130 K-gluconate, 4 MgATP, 0.3 NaGTP, 10 HEPES, 0.5 EGTA, 5 KCl, 10 Na_2_-phosphocreatine (internal solution 2, 291 mOsm, pH was adjusted to 7.3 with KOH). The recordings with either solution 1 or 2 produced similar results, and we combined the data from both experimental sets.

*The resting membrane potential* (*RMP)* was measured within 1-2 minutes after seal breaking. Once the membrane potential was stable, 3-10 seconds of the membrane potential was averaged and determined as the RMP.

To calculate *input resistance (R_in_)*, the neuron was held approximately at −60 mV, and a hyperpolarizing step of −10 pA for 400 ms was given. R_in_ was calculated by dividing the voltage response at the steady state by the current injection amplitude. At least 2 repetitions were used and averaged to calculate R_in_.

For the *f-I* (frequency-intensity) curve, PNs were held approximately at −70 mV, and a 500 ms current step was given at 20 pA increments every 10 seconds. Firing type was classified by the following criteria^61^: (1) tonically firing cells showed AP discharge throughout the current injection; (2) delayed firing cells had a delayed onset to the first AP and a hyperpolarizing notch after the initial depolarization at the start of the current injection; (3) phasically firing cells ceased AP firing in the second half of the current injection or showed only an initial burst of APs; (4) H-current cells showed a sag potential that peaked at ∼55 ms following termination of the current injection; (5) Cells that could not be classified by one of the criteria above, were named unidentified. Only the properties of PNs exhibiting delayed firing patterns were analyzed.

##### Action potential (AP) frequency

For all Figures except Figs. 4c and 4f, frequencies were calculated by dividing the number of APs by the duration of the current injection (500 ms). For Figs. 4c and 4f, frequencies were calculated by dividing the number of APs by the time interval between the time point of the first AP threshold and the end of the current injection. *f-I curve slope* was calculated by extracting the slope of the linear fit for the firing frequencies starting in the pulse preceding the first action potential firing.

*Latency to the first AP* was calculated as the time interval between the beginning of the current injection and the time point of the first AP threshold.

To calculate *AP threshold, half-width, and after-hyperpolarization (AHP),* the first spike of the first suprathreshold voltage response in the *f-I* protocol was analyzed. *AP threshold* was calculated using a phase plot analysis as previously described^62^. The point in the phase plot where the increase in dV/dt was larger than 8 mV/ms was determined as the AP threshold. *AP half-width* was determined as the width at half amplitude between the AP threshold and peak.

*AP AHP* was determined as the difference between the AP threshold and the minimal membrane potential 5-10 ms after the AP peak.

*AP amplitude* and *dV/dt_max_* were calculated for APs evoked by a short, 10 ms, current steps increasing by 10 pA increments from a holding potential of approximately −70 mV. The AP of the first suprathreshold voltage response was analyzed. The AP Amplitude was determined as the amplitude between the AP threshold and peak. *dV/dt_max_* was determined as the maximal point of phase plot analysis.

Recordings of AP firing before and after application of 5 mM 4-aminopyridine (4-AP) were performed in the presence of 10 µM NBQX to block excitatory postsynaptic potentials (EPSPs). The recordings were performed at least 5 minutes following the perfusion of 4-AP.

#### Voltage-clamp recordings

For voltage-clamp recordings, data were sampled at 50 kHz and low pass filtered at 2 kHz. Recordings commenced at least 3 minutes after the seal broke. Capacitive transient and leak currents were not compensated, and recordings were discarded if R_s_ was >20 MΩ or changed >15% during recordings.

For I_A_ recordings, 0.5 µM tetrodotoxin (TTX), 100 µM CdCl_2_, 10 µM NBQX, and 100 µM picrotoxin were added to bath solution to block voltage-gated Na^+^ currents, Ca^2+^ currents, excitatory and inhibitory postsynaptic currents (EPSCs and IPSCs), respectively. The internal solution contained (in mM): 130 K-gluconate, 4 MgATP, 0.3 NaGTP, 10 HEPES, 0.5 EGTA, 5 KCl, 10 Na_2_-phosphocreatine (291 mOsm, pH was adjusted to 7.3 with KOH). *To isolate I_A_*, membrane voltage was held at –80 mV in voltage-clamp mode. A two-step voltage protocol was used to determine the voltage-dependence of activation with a series of 500 ms depolarizing voltage steps in +10 mV increments to a maximum of +40 mV at 8-second intervals (**Supplementary Fig. 7a**, *left*). Next, a corresponding paired voltage step was given, consisting of the same voltage command but with a 150 ms prepulse to −10 mV (**Supplementary Fig. 7a**, *right*). I_A_ was extracted by the digital subtraction of each pair of pulses. I_A_ amplitudes were calculated from the isolated I_A_ traces as the current difference between the minimal current after the transient capacitive current and the peak of the current.

##### Voltage-dependence of I_A_ activation

I_A_ amplitudes were converted to conductances using the equation g = I/(V_m_ – V_rev_) where V_m_ is the membrane voltage at the voltage step, and V_rev_ is the calculated K^+^ reversal potential (−88.85 mV). Normalized conductances were fitted with a Boltzmann equation: g/g_max_ = 1/(1+exp((V_1/2_ – V) /*k*)), where g_max_ is the maximal conductance, V_1/2_ is the half-maximal voltage and *k* is the slope factor.

*Voltage-dependence of I_A_ inactivation* was determined using incremental conditioning command of 150 ms prepulses from −100 mV to +20 mV, followed by a voltage step to +40 mV for 500 ms at 8-second intervals. All traces were subtracted with the trace obtained at the final voltage step to zero residual currents. Current amplitudes were calculated as the difference between the peak amplitude and the mean steady-state current in the last 20 ms of the voltage command. Normalized currents were fitted similarly with the Boltzmann equation: I/I_max_ = 1/(1+exp((V_1/2_ – V) /*k*)), where I_max_ is the maximal current, V_1/2_ is the half-maximal voltage and *k* is the slope factor.

*I_A_ window current* was calculated as the area under the activation and inactivation curves for each cell.

I_A_ kinetics were quantified for the isolated I_A_ traces at the peak conductance (+10 mV). Decay current was fitted with 1 or 2 exponents, depending on the fit with the minimal residual sum of squares between the data and the fit. For most cells, 2 exponents resulted in the best fit (Control: 7/9; Inflammation: 6/10; Sham: 8/14; CCI-ION: 5/7).

All electrophysiological data were analyzed using custom scripts in Matlab (MathWorks, Natick, MA, USA).

### Immunofluorescence staining

Mice were deeply anesthetized with isoflurane and perfused transcardially with 20 ml of cold phosphate-buffered saline (PBS) followed by 20 ml of cold 4% paraformaldehyde (PFA). Trigeminal ganglions (TGs) and brains were extracted and post-fixed in 4% PFA 2 hours and overnight, respectively, at 4°C. Next, tissues were rinsed in PBS (3×10 min), followed by cryoprotection in 30% (w/v) sucrose in PBS for at least 2 nights at 4°C. Next, brains and TGs were frozen in OCT for sectioning. For TG staining, 15 µm free-floating TG sections were washed (3×10 min) in PBS. Following the initial washing steps, the sections were blocked with CAS-Block universal blocking agent (#008120, ThermoFisher Scientific) for 10 min at room temperature. The sections were then incubated in goat anti-c-Fos primary antibody (1:500, #sc-52-G, Santa Cruz Biotechnology) diluted in the same blocking buffer overnight at 4°C. On the next day, the slices were washed (3×10 min) in PBS and incubated with Alexa Fluor 488-conjugated donkey anti-goat secondary antibodies (1:1000, #705545147, Jackson ImmunoResearch) diluted in freshly prepared PBS in 0.03% Triton X-100 (#X100, Sigma Aldrich) for 2 hours at room temperature. Final wash steps were performed (3×10 min) in PBS. All the above steps were carried out on a shaker with gentle shaking at 70 rpm. Finally, the sections were mounted onto the microscope slides and air-dried before covering them with VECTASHIELD antifade mounting media with DAPI (#EW9395224, Vector Laboratories). The slices were then coverslipped for imaging. Imaging was done with a confocal microscope (Nikon Ti Eclipse) using a 20X objective at a resolution of 2048 pixels. cFos expressing cells were identified using custom scripts in Cellprofiler^63^ and validated by eye. For medullary dorsal horn staining, 30 µm free-floating sections (3-5 slices for each mouse) were washed in PBS and blocked in a solution containing 10% normal donkey serum and 0.3% Triton X-100 in PBS for 1 hour at room temperature. Rabbit anti-c-Fos primary antibody (1:500, #226 003, Synaptic Systems) was diluted in a solution containing 5% normal donkey serum and 0.3% Triton X-100 in PBS and incubated with sections overnight at 4°C. The following day, sections were rinsed in PBS (3×10 min) and incubated with Donkey anti-Rabbit Alexa Fluor 647 (1:1000, ab150063, abcam) secondary antibody diluted in a solution containing 5% normal donkey serum and 0.3% Triton X-100 in PBS for 2 hours. Next, slices were incubated in DAPI (2 µg/ml, 10236276001, Roche Merck) in PBS (10 min), followed by two additional washes in PBS (2×10 min). Finally, slices were mounted (Dako fluorescence mounting medium) and coverslipped. Slices were imaged using Olympus IX83P2ZF with a 10x objective lens. For analysis, the medullary dorsal horn region was cropped for each slice using ImageJ. Using Cellprofiler custom scripts, c-Fos and tdTomato expressing cells were identified and validated by eye. For each mouse, the percentage of co-labeled cells was calculated as the number of c-Fos+ and tdTomato expressing cells from all slices out of the total number of tdTomato expressing cells. TG and medullary dorsal horn c-Fos stainings were done by an experimenter blinded to mice groups.

### Computational modeling

Projection neurons were simulated using NEURON simulation environment^64^. Neurons were approximated to a single-compartment cylindrical model with both diameter and length of 45.75 *μm*. This was chosen in order to achieve the measured average capacitance of 65.8 *pF* (**Supplementary Fig. 3**), with a standard specific capacitance of 1 *μF*/*cm*^2^.

All active and passive mechanisms were adapted from Medlock et al. 2022^38^. For the *I*_*A*_ mechanism, the inactivation parameter *l*, was altered to follow a double exponential curve rather than the standard single exponential to fit our experimental measurements.

#### Passive properties

The passive conductance value (*g*_*pas*_) was determined according to the measured input resistance (*R*_*in*_ = 614 *M*Ω, **Fig. 1i**) and was equal to *g*_*pas*_ = 1.372 × 10^−5^ *S*/*cm*^2^. The equilibrium potential of the passive mechanism was set at −58 *mV* to achieve a resting membrane potential of −65 *mV*, which is within the range of our measurements in control mice (**Fig. 1j**).

The potassium reversal potential (*E*_*k*_) was set at −88.85 *mV* in accordance with the solutions we used in the experiments. The sodium reversal potential (*E*_*Na*_) was kept at the standard 50 *mV*.

#### Active properties

Three active conductance mechanisms were included in the model: (1) a Hodgkin-Huxley mechanism, including both a sodium conductance 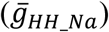 and a potassium conductance 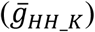. (2) A delayed rectifier K^+^ (KDR) mechanism with a single potassium conductance 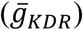, and (3) an *I*_*A*_ mechanism with a single potassium conductance 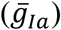. The values assigned to these parameters were:

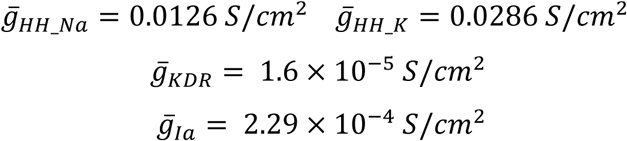

In addition to these parameters, the *I*_*A*_ mechanism included parameters corresponding to the activation and inactivation kinetics of the potassium channel given by the mechanism state variables *n* for the activation and *l* for the inactivation, such that the *I*_*A*_ conductance is given by:

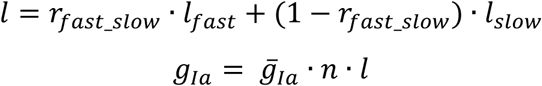

And the evolution of *n* and *l* were governed by the equations:

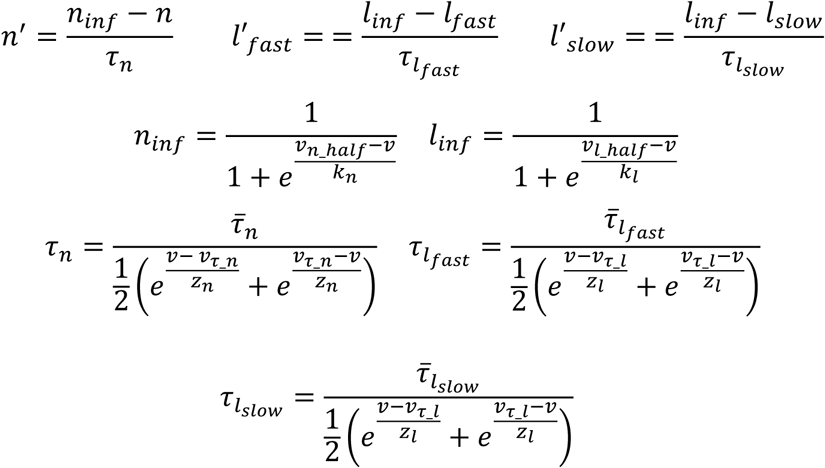

where *v* is the membrane voltage.

The base parameters used in the simulation of PNs in control conditions are:

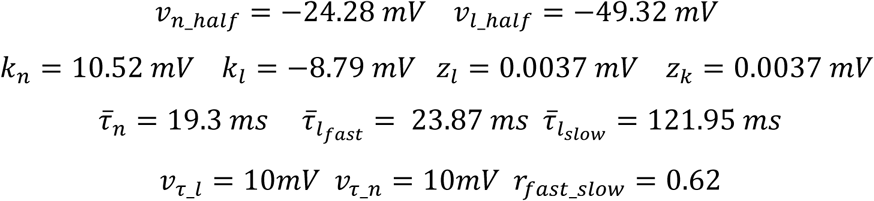

The parameters 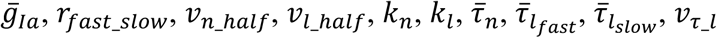, and *v*_*τ*_*n*_ (for which we had experimental measurements) were chosen as the average measured values of these parameters in control conditions. The additional model parameters were given their specific values in order to replicate the observed firing pattern of the cells from the control group of our experiments.

The cell was held at −70 *mV* for 500 *ms* before injecting a constant current ranging between 60 − 140 *pA* with 20 *pA* intervals for 500 *ms*. Latencies to the first AP and firing frequencies were calculated for increasing values of the *v*_*l*_ℎ*alf*_ parameter, starting at the initial value specified above and steadily increasing in 5 constant intervals to the average value observed in the acute pain conditions, which was −45.2079 *mv*.

All model parameters and the complete code used for simulation are available on Git Hub.

### Statistics

Experiments were performed with side-by-side control and in a random order. No power analysis was used to determine experimental sample sizes in advance, but sample sizes were similar to those reported previously. The number of mice, slices, and cells are indicated in the Supplementary Tables accompanying each figure. Parametric and non-parametric tests were used where appropriate. All statistical tests, statistics (*F* or *t*), and *P* values are reported in figures legends or in the Supplementary Tables accompanying each figure. For the statistical analysis of eye wipes over time (**Fig. 1f**, *left*) we fitted a linear mixed-effects model (MATLAB, function *fitlme*) using *group, day,* and the interaction of them for fixed effects and *mouse id* for a random effect. For the statistical analysis of *f*-I curves, latency to 1^st^ AP, and intrinsic properties between 3 groups (**Fig. 2**), we fitted a linear mixed-effects model using *group* or *condition* (**Fig. 5a**) for a fixed effect. In repeated measurements structured experiments (*f*-I curves and latency to 1^st^ AP), we usually used *mouse id*, *neuron id,* and *current injection* for random effects unless stated otherwise in the statistics table. For intrinsic properties analysis in **Fig. 2b,c**, we used *mouse id* for a random effect. The random effects were varied specifically for each case to improve the fit of the model to the data. This was examined using *compare* function in MATLAB. All models are depicted in the statistics table. All statistical analyses were conducted with Matlab (MathWorks, Natick, MA, USA).

## Acknowledgments

This work was funded by: The Israel Science Foundation – Individual research grant 1202/23 (AB), Israel Cancer Research Fund (ICRF) – The Brause Family Initiative for Quality of Life 22-402-QOL (AB), The Israel Science Foundation (ISF) and the Azrieli Foundation - 2545/18 (AB), and The Deutsch-Israelische Projectkooperation program of the Deutsche Forschungsgemeinschaft (DIP)- B.I. 1665/1-1ZI1172/12-1 (AB) and Cecile and Seymour Alpert Chair in Pain Research (AB).

## Author contributions

Conceptualization, B.T, Y.Y. and A.B.; Investigation, B.T, E.V., N.E., P.R.R., R.Y., S.H., S.L.; Formal Analysis, B.T, E.V., N.E., and A.B.; Writing – Original Draft, B.T, and A.B.; Writing – Review & Editing, B.T, B.K., S.L., and A.B.; Funding Acquisition, A.B.; Supervision, Y.Y., A.B.

## Compelling interests

The authors declare that they have no competing interests.

## Materials and correspondence

Correspondence should be sent to: **AMB** (Department of Medical Neurobiology; Institute for Medical Research Israel-Canada, The Hebrew University-Hadassah School of Medicine, POB 12271, Jerusalem, Israel, 91120). All datasets generated and/or analyzed during the current study are available in the main text or upon request. All model parameters and the complete code used for simulation are available on Git Hub.

Further information and requests for resources and reagents should be directed to and will be fulfilled by the Lead Contact, Alexander Binshtok.

