## Supplementary Figures and legends for "Adaptation of pain-related projection neurons in acute but not chronic pain"

**Supplementary Information**

**Supplementary Figures and Legends**

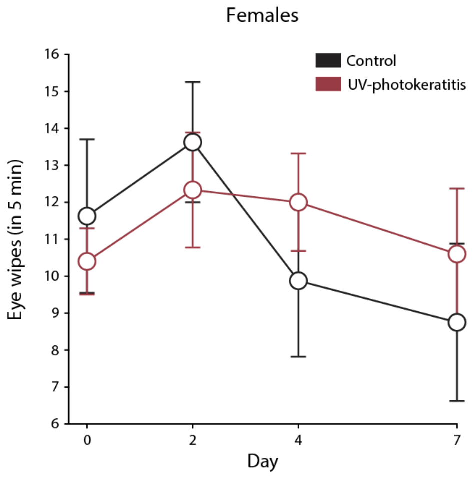

**Supplementary Fig. 1, related to Fig. 1**

Number of eye wipes (measured in 5 min) following capsaicin application to the eyes of female mice in acute pain (UV-photokeratitis, *red*) and control conditions (*black)*. Control: N = 8, UV-photokeratitis: N = 12. Data are presented as mean ± s.e.m.

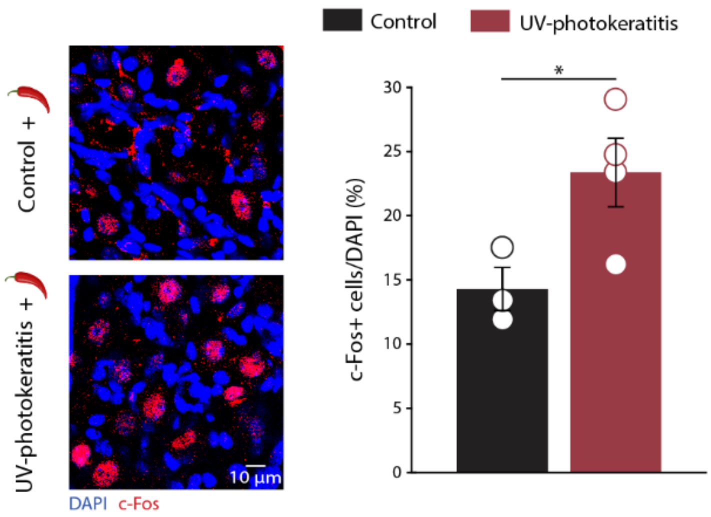

**Supplementary Fig. 2, related to Fig. 1**

*Left,* Representative images of trigeminal ganglion neurons stained for c-Fos following capsaicin application to the eye from mice in acute pain and control conditions. *Right,* Bar graph of means ± s.e.m and values from individual mice of activated TG neurons (TG neurons expressing c-Fos, normalized to DAPI) following capsaicin application to the eye from mice in acute pain *(red)* and control conditions (*black*). Each data point represents the mean percentage of activated TG neurons measured from individual mice. Control: N = 3; UV-photokeratitis: N = 4, $P = 0.047$. Significance was assessed by a two-tailed unpaired student's *t*-test. * $P < 0.05$. Please refer to Supplementary Table 1 for full statistical information.

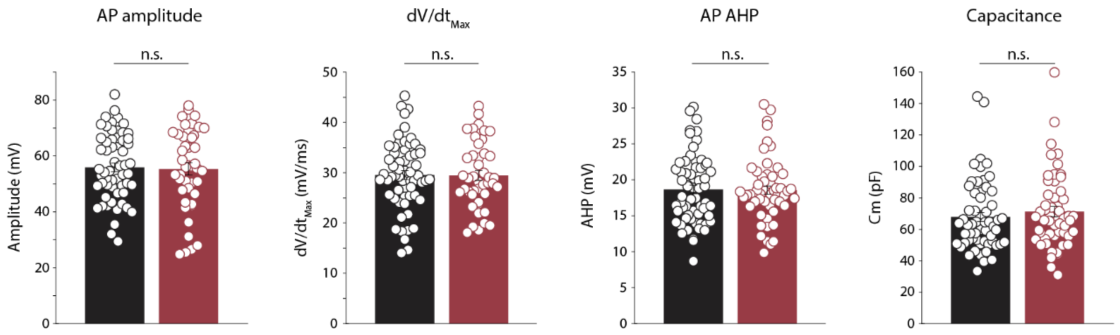

**Supplementary Fig. 3, related to Fig. 1**

Bar graphs of means ± s.e.m and individual values of the intrinsic properties (specified above the panel) of PNs from mice in acute pain (*red)* and control conditions (*black)*. *AP amplitude*, Control {n cells/N mice}: 61/14, UV-photokeratitis: 44/10, *P* = 0.8217; *dV/dt$_{Max}$*, Control: 61/14, UV-photokeratitis: 44/10, *P* = 0.937; *AP after-hyperpolarization*, Control: 62/14, UV-photokeratitis 55/13, *P* = 0.9164; *Capacitance*, Control: 59/14, UV-photokeratitis: 54/13, *P* = 0.438. Significance was assessed by a two-tailed unpaired student's *t*-test. n.s. not significant. Please refer to Supplementary Table 1 for full statistical information.

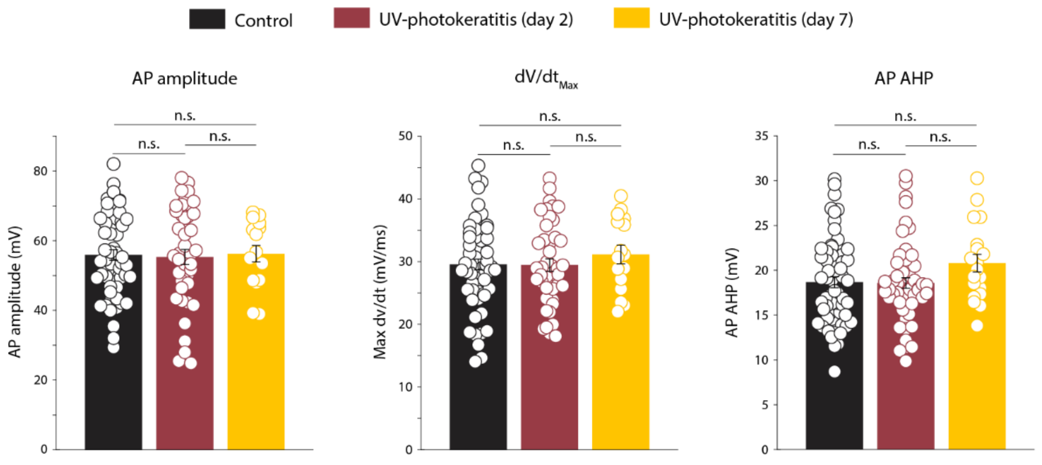

**Supplementary Fig. 4, related to Fig. 2**

Bar graphs of means ± s.e.m and individual values of the intrinsic properties (specified above the panel) of PNs from control mice (*black*) and mice 2 days (acute pain, *red*) and 7 days (recovery, *yellow*) after UV irradiation. *AP amplitude*, Control {n cells/N mice}: 61/14, Inflammation (day 2): 44/10, UV-photokeratitis (day 7): 16/4, $P = 0.9593$; $dV/dt_{Max}$, Control: 61/14, UV-photokeratitis (day 2): 44/10, UV-photokeratitis (day 7): 16/4, $P = 0.6557$; *AP after-hyperpolarization*, Control: 62/14, UV-photokeratitis 55/13, UV-photokeratitis (day 7): 19/4, $P = 0.1608$. Significance was assessed by one-way ANOVA. n.s. not significant. Please refer to Supplementary Table 2 for full statistical information.

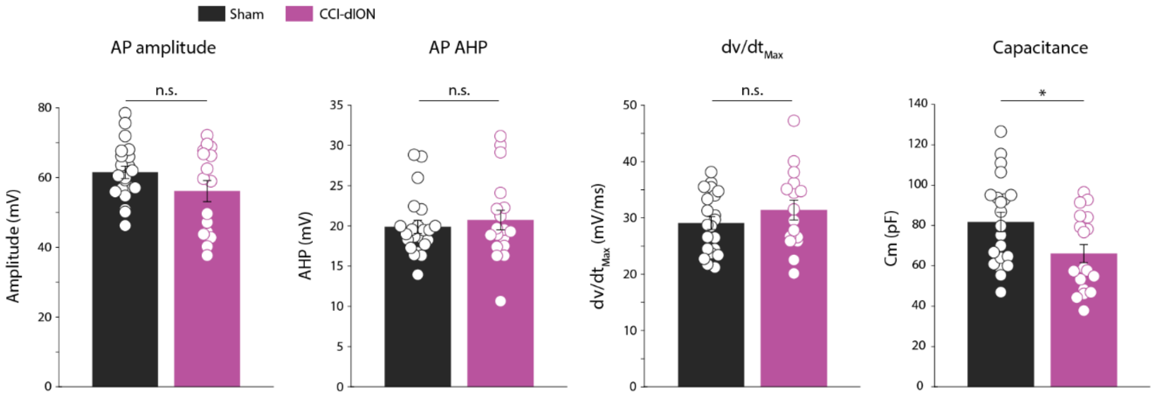

**Supplementary Fig. 5, related to Fig. 3**

Bar graphs of means ± s.e.m and individual values of the intrinsic properties (specified above the panel) of PNs from mice in chronic pain (2-4 weeks after CCI-dION, *purple*) and sham conditions (*black*). *AP amplitude*, Sham 21/5, CCI-dION 16/4, $P$ = 0.1171; $dV/dt_{Max}$, Sham: 21/5, CCI-dION: 16/4, $P$ = 0.2644; *AP after-hyperpolarization*, Sham 22/5, CCI-dION 18/4, $P$ = 0.5575). n.s. not significant. Please refer to Supplementary Table 3 for full statistical information.

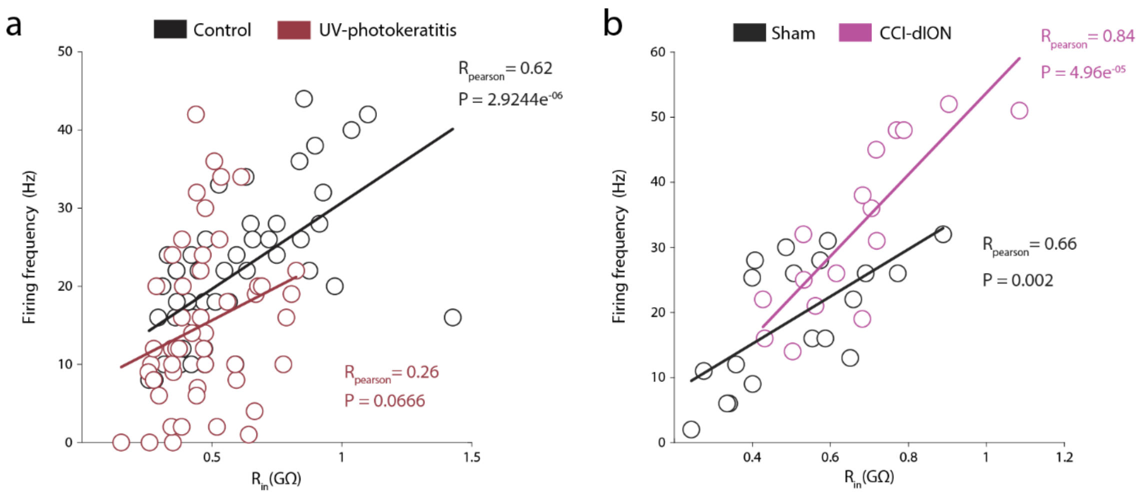

**Supplementary Fig. 6, related to Fig. 4.**

**a,** Correlation between input resistance and firing frequency (measured at 140 pA current pulse) in PNs from mice in acute pain (*red)* and control conditions (*black)*. Control {n cells/N mice}: 48/24, UV-photokeratitis: 49/19. Solid lines represent the linear regression of the data points.

**b**, same as *a* but showing the correlation between input resistance and firing frequency (measured at 140 pA current pulse) in PNs from mice in chronic pain (*purple)* and control conditions (*black)*.

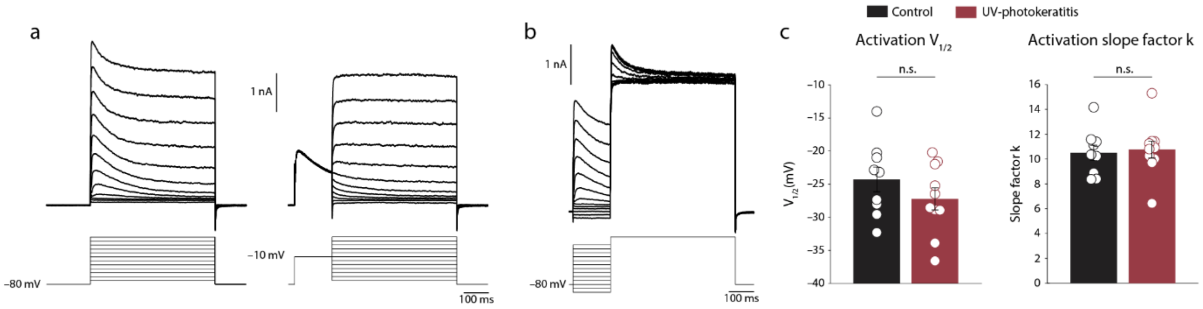

**Supplementary Fig. 7, related to Fig. 5**

**a,** *Left,* Representative traces of voltage clamp recordings of PN's responses to a series of increasing depolarizing voltage steps of +10 mV to isolate $I_A$. *Right,* a protocol used to assess the voltage dependence of $I_A$ activation. The responses of the neuron shown in the left panel to a similar series of depolarizing voltage steps but with a prepulse command to −10 mV for 150 ms to inactivate the transient currents.

**b,** a protocol used to assess the voltage dependence of $I_A$ inactivation. Representative traces of PN responses to a voltage step to +40 mV following a series of depolarizing prepulses from −100 mV to +20 mV.

**c,** Bar graphs of means ± s.e.m and individual values of $V_{1/2}$ (*left*) and slope factor k (*right)* of the $I_A$ activation curve in acute pain (*red)* and control conditions (*black*). Control: 9/7, UV-photokeratitis: 10/4, $V_{1/2}$: $P = 0.255$; slope factor k: $P = 0.7883$.

Significance was assessed by a two-tailed unpaired student's *t*-test. n.s. not significant. Please refer to Supplementary Table 5 for full statistical information.

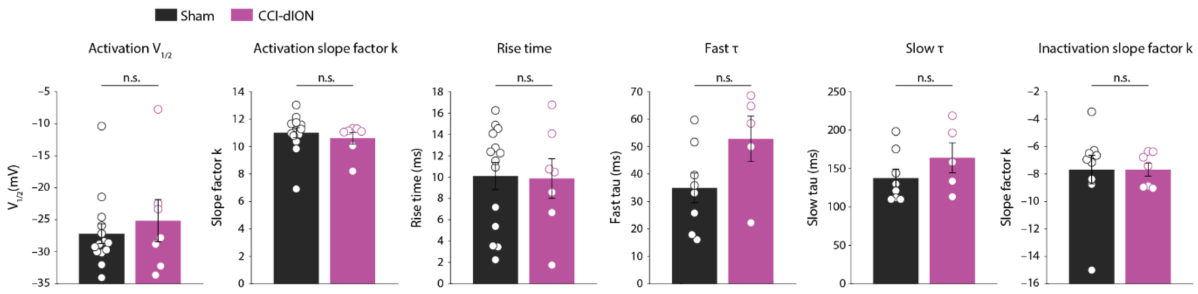

**Supplementary Fig. 8, related to Fig. 5.**

Comparison of $I_A$ properties (indicated above the panels, means ± s.e.m and individual values) from PNs in chronic pain (*purple)* and sham conditions (*black*). *$V_{1/2}$ of $I_A$ activation and slope factor k*, Sham: 14/5, CCI-dION: 7/5, $V_{1/2}$: $P$ = 0.5286; slope factor k: $P$ = 0.525; *rise time*, Sham: 14/5, CCI-dION: 7/5, $P$ = 0.9187; *fast and slow τ*, Sham: 8/5, CCI-dION: 5/5, *fast τ*: $P$ = 0.0852; *slow τ*, $P$ = 0.2377). *$I_A$ inaactivation slope k*, Sham: 9/5, CCI-dION: 7/5, $P$ = 0.9977. Significance was assessed by two-tailed unpaired student's *t*-test. n.s. not significant. Please refer to Supplementary Table 5 for full statistical information.

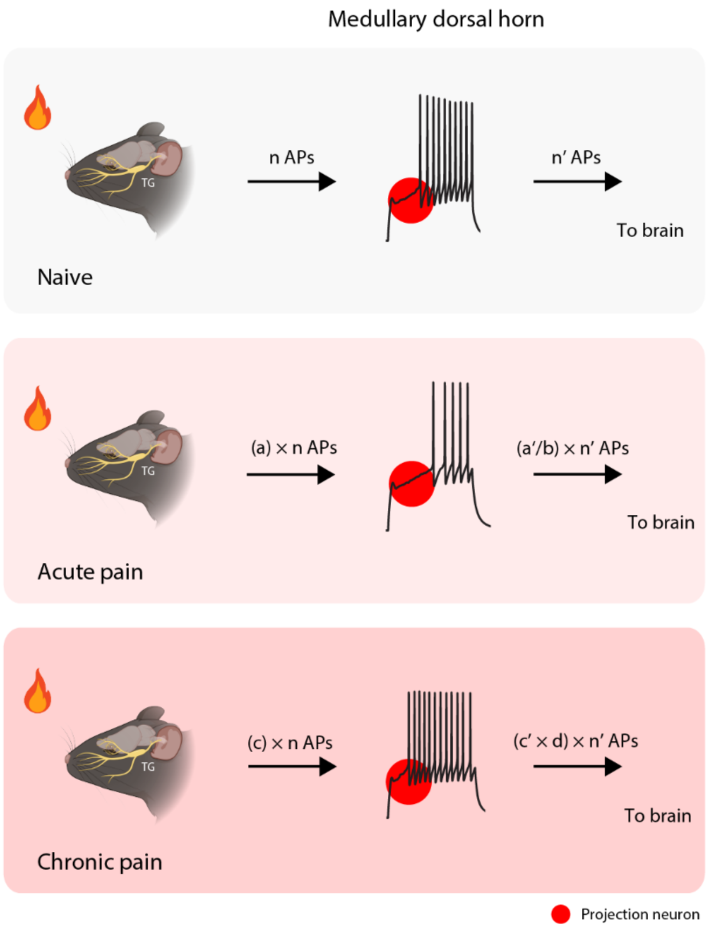

**Supplementary Fig. 9**

A scheme depicting PNs adaptation and its effect on the dorsal horn output.

*Upper,* In naïve conditions, a noxious stimulus activates peripheral neurons, triggering the firing of "*n*" APs. PNs summate the inputs from peripheral neurons and local interneurons and fire a proportional number of APs ("*n'*" APs).

*Middle,* In acute pain conditions, the combined effect of an increased overall number of activated peripheral neurons (**Supplementary Fig. 2**) and an increase in the excitability amplify (by a factor *"a"*) the output from the peripheral neurons towards the medullary dorsal horn (*"a * n"* APs). This increased activity would both directly (via monosynaptic connections) and indirectly (by increased activity of local interneurons[20,65]) amplify the firing towards the PNs ("*a'*n*"). However, the PN adaptation in acute pain conditions, which leads to a decrease in their excitability (by a factor "*b*"), tunes down PNs output towards higher centers ("*a'/b * n'*" APs).

*Bottom,* In chronic pain conditions, the output from peripheral neurons also increases[33], but this time by a factor "*c*." However, in these conditions, increased PNs excitability (factor "*d*") further amplifies the enhanced input from the peripheral neurons and local interneurons, leading to increased output from PNs towards higher brain centers, plausibly triggering and facilitating pain chronification.

**Supplementary Table 1, Statistical information for Figure 1 and Supplementary Figures 1-3**

| Figure | N | Statistical test | Statistic | P. value |
|---|---|---|---|---|
| 1e *left* | 6 mice per group | Two-way ANOVA for repeated measures with Tukey's multiple comparisons test | Group: $F_{1,10}$ = 9.7328<br>Group × days: $F_{1,10}$ = 5.7669 | Group: $P$ = 0.010879<br>Group × days: $P$ = 0.037225<br>Post-hoc Tukey's test for day 2: $P$ = 0.0072708 |
| 1e *right* | 6 mice per group | Two-tailed unpaired student's *t*-test | $t_{10}$ = 2.7538 | $P$ = 0.0203 |
| 1f *left* | Control: N = 7, UV-photokeratitis: N = 9 | Linear mixed effects model: $Eye\ wipes \sim Group \times Day + (1\|MouseID)$ | Group: $F_{1,101}$ = 3.9164<br>Group × days: $F_{1,101}$ = 2.2061 | Group: $P$ = 0.050538<br>Group × days: $P$ = 0.14058 |
| 1f *right* | Control: N = 7, UV-photokeratitis: N = 9 | Two-tailed unpaired student's *t*-test | $t_{14}$ = 2.7538 | $P$ = 0.0484 |
| 1g *right* | Control {n cells/N mice}: 60/26, UV-photokeratitis: 51/19 | Linear mixed effects model: $Frequency \sim Group + (1\|NeuronID) + (1\|MouseID) + (Group\|CurrentInjectior$ | $F_{1,759}$ = 3.9164 | $P$ = 0.014744 |
| 1h | Control: 61/26, UV-photokeratitis: 51/20 | Two-tailed unpaired student's *t*-test | $t_{110}$ = 2.2513 | $P$ = 0.0264 |
| 1i *right* | Control: 59/26, UV-photokeratitis: 54/20 | Two-tailed unpaired student's *t*-test | $t_{111}$ = 3.1114 | $P$ = 0.0024 |
| 1j *AP half-width* | Control: 62/26, UV-photokeratitis: 55/20 | Two-tailed unpaired student's *t*-test | $t_{115}$ = -2.5516 | $P$ = 0.012 |
| 1j *AP threshold* | Control: 62/26, UV-photokeratitis: 55/20 | Two-tailed unpaired student's *t*-test | $t_{115}$ = -1.495 | $P$ = 0.1377 |
| 1j *RMP* | Control: 56/26, UV-photokeratitis: 49/20 | Two-tailed unpaired student's *t*-test | $t_{103}$ = 1.4973 | $P$ = 0.1374 |
| Supplementary fig. 1 | Control: N = 9, UV-photokeratitis: N = 13 | Linear mixed effects model: $Eye\ wipes \sim Group \times Day + (1\|MouseID)$ | Group: $F_{1,68}$ = 2.6084<br>Group × days: $F_{1,68}$ = 2.8679 | Group: $P$ = 0.11093<br>Group × days: $P$ = 0.094942 |

| | | | | |
|---|---|---|---|---|
| Supplementary fig. 2 | Control: N = 6, UV-photokeratitis: N = 7, 3-5 slices per mouse | Two-tailed unpaired student's *t*-test | $t_5 = -2.6226$ | $P = 0.047$ |
| Supplementary fig. 3 *AP amplitude* | Control: 61/26, UV-photokeratitis: 44/20 | Two-tailed unpaired student's *t*-test | $t_{102} = 0.0578$ | $P = 0.954$ |
| Supplementary fig. 3 $dV/dt_{Max}$ | Control: 61/26, UV-photokeratitis: 44/20 | Two-tailed unpaired student's *t*-test | $t_{103} = 0.0793$ | $P = 0.937$ |
| Supplementary fig. 3 *AP AHP* | Control: 62/26, UV-photokeratitis: 55/20 | Two-tailed unpaired student's *t*-test | $t_{115} = 0.1052$ | $P = 0.9164$ |
| Supplementary fig. 3 *Capacitance* | Control: 59/26, UV-photokeratitis: 54/20 | Two-tailed unpaired student's *t*-test | $t_{111} = -0.7784$ | $P = 0.438$ |

**Supplementary Table 2, Statistical information for Figure 2, and Supplementary Figure 4**

| Figure | N | Statistical test | Statistic | P. value |
|---|---|---|---|---|
| 2a *right* | Control {n cells/N mice}: 60/26, UV-photokeratitis (day 2): 51/19, UV-photokeratitis (day 7): 19/4 | Linear mixed effects model: $Frequency \sim Group + (1\|NeuronID) + (1\|MouseID) + (Group\|CurrentInjectio$ | $F_{2,889}$ = 3.545 | $P$ = 0.029276 Control vs UV: $P$ = 0.0149, Control vs recovery: $P$ = 0.3097, UV vs recovery: 0.0216 |
| 2b | Control: 61/26, UV-photokeratitis (day 2): 51/20, UV-photokeratitis (day 7): 19/4 | Linear mixed effects model: $Slope \sim Group + (1\|MouseID)$ | $F_{2,128}$ = 4.805 | $P$ = 0.0097241 Control vs UV: $P$ = 0.039, Control vs recovery: $P$ = 0.1237, UV vs recovery: 0.0045 |
| 2c *right* | Control: 59/26, UV-photokeratitis (day 2): 49/20, UV-photokeratitis (day 7): 19/4 | Linear mixed effects model: $Rin \sim Group + (1\|MouseID)$ | $F_{2,129}$ = 5.9474 | $P$ = 0.0033831 Control vs UV: $P$ = 0.0031, Control vs recovery: $P$ = 0.5015, UV vs recovery: $P$ = 0.0099 |
| 2d *AP half-width* | Control: 62/26, UV-photokeratitis (day 2): 55/20, UV-photokeratitis (day 7): 19/4 | Linear mixed effects model: $HalfWidth \sim Group + (1\|MouseID)$ | $F_{2,133}$ = 3.0771 | $P$ = 0.049391 Control vs UV: $P$ = 0.1663, Control vs recovery: $P$ = 0.1087, UV vs recovery: $P$ = 0.0182 |
| 2d *AP threshold* | Control: 62/26, UV-photokeratitis (day 2): 55/20, UV-photokeratitis (day 7): 19/4 | Linear mixed effects model: $Threshold \sim Group + (1\|MouseID)$ | $F_{2,133}$ = 1.0431 | $P$ = 0.35523 |
| 2d *RMP* | Control: 56/26, | Linear mixed effects model: | $F_{2,120}$ = 11.619 | $P = 2.4391e^{-05}$ Control vs UV: |

| | | | | |
|---|---|---|---|---|
| | UV-photokeratitis (day 2): 49/20, UV-photokeratitis (day 7): 18/4 | $RMP \sim Group + (1\|MouseID)$ | | $P = 0.2206$, Control vs recovery: $P = 4.2502e^{-06}$, UV vs recovery: $P = 1.1462e^{-04}$ |
| Supplementary fig. 4 *AP amplitude* | Control: 61/26, UV-photokeratitis (day 2): 44/20, UV-photokeratitis (day 7): 16/4 | One-way ANOVA with Tukey's multiple comparisons test | $F_{2,118} = 0.0416$ | $P = 0.9593$ |
| Supplementary fig. 4 $dV/dt_{Max}$ | Control: 61/26, UV-photokeratitis (day 2): 44/20, UV-photokeratitis (day 7): 16/4 | One-way ANOVA with Tukey's multiple comparisons test | $F_{2,118} = 0.4236$ | $P = 0.6557$ |
| Supplementary fig. 4 *AP AHP* | Control: 62/26, UV-photokeratitis (day 2): 55/20, UV-photokeratitis (day 7): 19/4 | One-way ANOVA with Tukey's multiple comparisons test | $F_{2,133} = 1.8532$ | $P = 0.1608$ |

**Supplementary Table 3, Statistical information for Figure 3, and Supplementary Figure 5**

| Figure | N | Statistical test | Statistic | P. value |
|---|---|---|---|---|
| 3b | 7 mice per group | 2-way ANOVA for repeated measures with Tukey's multiple comparisons test | Group: $F_{1,12}$ = 70.877 Group × time: $F_{1,12}$ = 114.72 | Group: $P$ = 2.2214e$^{-06}$ Group × time: $P$ = 1.6972e$^{-07}$ |
| 3c | 6 mice per group | Two-tailed unpaired student's *t*-test | $t_{10}$ = 4.7867 | $P$ = 7.382e$^{-04}$ |
| 3d *right* | Sham {n cells/N mice}: 22/4, CCI-dION: 18/4 | Linear mixed effects model: $Frequency \sim Group + (Group \mid NeuronID) + (1 \mid CurrentInjection)$ | $F_{1,274}$ = 8.8714 | $P$ = 0.0031561 |
| 3e | Sham: 22/4, CCI-dION: 18/4 | Two-tailed unpaired student's *t*-test | $t_{38}$ = -2.6135 | $P$ = 0.0128 |
| 3f *right* | Sham: 21/4, CCI-dION: 18/4 | Two-tailed unpaired student's *t*-test | $t_{37}$ = -2.4227 | $P$ = 0.0204 |
| 3g *AP half-width* | Sham: 22/4, CCI-dION: 18/4 | Two-tailed unpaired student's *t*-test | $t_{38}$ = 3.4752 | $P$ = 0.0013 |
| 3g *AP threshold* | Sham: 22/4, CCI-dION: 18/4 | Two-tailed unpaired student's *t*-test | $t_{38}$ = -0.6248 | $P$ = 0.5358 |

| | | | | |
|---|---|---|---|---|
| 3g *RMP* | Sham: 21/4, CCI-dION: 18/4 | Two-tailed unpaired student's *t*-test | $t_{37}$ = -0.5239 | *P* = 0.6035 |
| Supplementary fig. 5 *AP amplitude* | Sham: 21/4, CCI-dION: 16/4 | Two-tailed unpaired student's *t*-test | $t_{35}$ = 1.6068 | *P* = 0.1171 |
| Supplementary fig. 5 *dV/dt<sub>Max</sub>* | Sham: 21/4, CCI-dION: 16/4 | Two-tailed unpaired student's *t*-test | $t_{35}$ = -1.1342 | *P* = 0.2644 |
| Supplementary fig. 5 *AP AHP* | Sham: 22/4, CCI-dION: 18/4 | Two-tailed unpaired student's *t*-test | $t_{38}$ = -0.5918 | *P* = 0.5575 |
| Supplementary fig. 5 *Capacitance* | Sham: 21/4, CCI-dION: 18/4 | Two-tailed unpaired student's *t*-test | $t_{37}$ = 2.3525 | *P* = 0.0241 |

**Supplementary Table 4, Statistical information for Figure 4, and Supplementary Figure 6**

| Figure | N | Statistical test | Statistic | P. value |
|---|---|---|---|---|
| 4a | Control {n cells/N mice}: 42/22, UV-photokeratitis: 38/19 | Pearson correlation | Control: $R_{Pearson}$ = -0.6871 UV-photokeratitis: $R_{Pearson}$ = -0.7375 | Control: $P$ = 5.0379e$^{-07}$ UV-photokeratitis: $P$ = 1.2762e$^{-07}$ |
| 4b | Control: 60/26, UV-photokeratitis: 47/19 | Linear mixed effects model: $Latency \sim Group + (1\|NeuronID) + (Group\|CurrentInjection$ | $F_{1,416}$ = 13.761 | $P$ = 0.00023576 |
| 4c | Control: 60/26, UV-photokeratitis: 51/19 | Linear mixed effects model: $Latency \sim Group + (1\|mouseID) + (Group\|NeuronID) + (Group\|CurrentInjection$ | $F_{1,754}$ = 2.9301 | $P$ = 0.087352 |
| 4d | Sham: 17/4, CCI-dION: 11/3 | Pearson correlation | Sham: $R_{Pearson}$ = -0.8752 CCI-dION: $R_{Pearson}$ = -0.8783 | Sham: $P$ = 4.2262e$^{-06}$ CCI-dION: $P$ = 0.00037493 |
| 4e | Sham: 22/4, CCI-dION: 18/4 | Linear mixed effects model: $Latency \sim Group + (1\|NeuronID) + (Group\|CurrentInjection$ | $F_{1,157}$ = 7.5022 | $P$ = 0.0068754 |
| 4f | Sham: 22/4, CCI-dION: 18/4 | Linear mixed effects model: $Frequency \sim Group + (Group\|NeuronID) + (1\|CurrentInjection)$ | $F_{1,274}$ = 8.9735 | $P$ = 0.0029899 |
| Supplementary fig. 6a | Control: 48/24, UV-photokeratitis: 49/19 | Pearson correlation | Control: $R_{Pearson}$ = -0.6176 UV-photokeratitis: $R_{Pearson}$ = -0.2642 | Control: $P$ = 2.9244e$^{-06}$ UV-photokeratitis: $P$ = 0.0666 |
| Supplementary fig. 6b | Sham: 19/4, CCI-dION: 16/4 | Pearson correlation | Sham: $R_{Pearson}$ = 0.6559 CCI-dION: $R_{Pearson}$ = 0.8385 | Control: $P$ = 0.0023 CCI-dION: $P$ = 4.9616e$^{-05}$ |

**Supplementary Table 5, Statistical information for Figure 5, and Supplementary Figures 7-8**

| Figure | N | Statistical test | Statistic | P. value |
|---|---|---|---|---|
| 5a *right* | {n cells/N mice}: 6/2 | Linear mixed effects model: $Latency \sim Condition + (1\|NeuronID) + (1\|CurrentInjection)$ | $F_{1,60}$ = 137.81 | $P$ = 3.4976e$^{-17}$ |
| 5c $V_{1/2}$ | control: 9/7, UV-photokeratitis: 9/4 | Two-tailed unpaired student's *t*-test | $t_{16}$ = -2.2289 | $P$ = 0.0405 |
| 5c *Slope factor k* | control: 9/7, UV-photokeratitis: 9/4 | Two-tailed unpaired student's *t*-test | $t_{16}$ = -0.7509 | $P$ = 0.4636 |
| 5d right | control: 7/7, UV-photokeratitis: 8/4 | Two-tailed unpaired student's *t*-test | $t_{13}$ = -2.2685 | $P$ = 0.041 |
| 5f *Rise time* | control: 9/7, UV-photokeratitis: 10/4 | Two-tailed unpaired student's *t*-test with unequal variance | $t_{13.0775}$ = -2.9441 | $P$ = 0.0113 |
| 5f *Fast tau* | control: 7/7, UV-photokeratitis: 6/4 | Two-tailed unpaired student's *t*-test with unequal variance | $t_{6.3676}$ = -2.4247 | $P$ = 0.0492 |
| 5f *Slow tau* | control: 7/7, UV-photokeratitis: 6/4 | Two-tailed unpaired student's *t*-test with unequal variance | $t_{6.0488}$ = -1.4454 | $P$ = 0.1981 |
| 5g | Control: 9/7, UV-photokeratitis: 10/4 | Two-tailed unpaired student's *t*-test | $t_{17}$= -2.125 | $P$ = 0.0485 |
| 5i $V_{1/2}$ | Sham: 9/5, CCI-dION: 7/5 | Two-tailed unpaired student's *t*-test with unequal variance | $t_{7.458}$ = -2.2685 | $P$ = 0.7468 |
| 5i *Window current* | Sham: 9/5, CCI-dION: 7/5 | Two-tailed unpaired student's *t*-test | $t_{14}$ = 0.6624 | $P$ = 0.5185 |
| 5j | Sham: 14/5, CCI-dION: 7/5 | Two-tailed unpaired student's *t*-test | $t_{19}$ = 0.2345 | $P$ = 0.8171 |
| Supplementary fig. 7c | control: 9/7, UV- | Two-tailed unpaired student's *t*-test | $t_{17}$ = 1.1781 | $P$ = 0.255 |

| | | | | |
|---|---|---|---|---|
| *Activation $V_{1/2}$* | photokeratitis: 10/4 | | | |
| Supplementary fig. 7c *Activation slope factor k* | control: 9/7, UV-photokeratitis: 10/4 | Two-tailed unpaired student's *t*-test | $t_{17}$ = -0.2728 | *P* = 0.7883 |
| Supplementary fig. 8a *Activation $V_{1/2}$* | Sham: 14/5, CCI-dION: 7/5 | Two-tailed unpaired student's *t*-test | $t_{19}$ = -0.6417 | *P* = 0.5286 |
| Supplementary fig. 8a *Activation slope factor k* | Sham: 14/5, CCI-dION: 7/5 | Two-tailed unpaired student's *t*-test | $t_{19}$ = 0.6476 | *P* = 0.525 |
| Supplementary fig. 8b *Rise time* | Sham: 14/5, CCI-dION: 7/5 | Two-tailed unpaired student's *t*-test | $t_{19}$ = 0.1034 | *P* = 0.9187 |
| Supplementary fig. 8b *Fast tau* | Sham: 8/5, CCI-dION: 5/5 | Two-tailed unpaired student's *t*-test | $t_{11}$ = -1.8912 | *P* = 0.0852 |
| Supplementary fig. 8b *Slow tau* | Sham: 8/5, CCI-dION: 5/5 | Two-tailed unpaired student's *t*-test | $t_{11}$ = -1.2487 | *P* = 0.2377 |
| Supplementary fig. 8c | Sham: 9/5, CCI-dION: 7/5 | Two-tailed unpaired student's *t*-test | $t_{14}$ = 0.0029 | *P* = 0.9977 |

**Supplementary Table 6, Statistical information for Figure 7**

| Figure | N | Statistical test | Statistic | P. value |
|---|---|---|---|---|
| 7b | 6 mice per group, 4-5 slices per mouse | Two-tailed unpaired student's *t*-test | $t_{10} = -3.5159$ | $P = 0.0056$ |
| 7d | Control: N = 6, UV-photokeratitis: N = 7, 3-5 slices per mouse | Two-tailed unpaired student's *t*-test | $t_{11} = -2.2302$ | $P = 0.0475$ |
